# Conserved Herpesvirus Protein Kinases Target SAMHD1 to Facilitate Virus Replication

**DOI:** 10.1101/399063

**Authors:** Kun Zhang, Dong-Wen Lv, Renfeng Li

**Affiliations:** Philips Institute for Oral Health Research, School of Dentistry, Virginia Commonwealth University, Richmond, Virginia, 23298, USA; Department of Microbiology and Immunology, School of Medicine, Virginia Commonwealth University, Richmond, Virginia, 23298, USA; Massey Cancer Center, Virginia Commonwealth University, Richmond, Virginia, 23298, USA

**Author notes:** Corresponding author (RL).

**Keywords:** Epstein-Barr virus, Conserved herpesvirus protein kinases, SAMHD1, lytic replication, phosphorylation, restriction factor, dNTPase

## Abstract

To ensure a successful infection, herpesviruses have developed elegant strategies to counterbalance the host anti-viral responses. Sterile alpha motif and HD domain 1 (SAMHD1) was recently identified as an intrinsic restriction factor for a variety of viruses. Aside from human immunodeficiency virus 2 (HIV-2) and the related simian immunodeficiency virus (SIV) Vpx proteins, the direct viral countermeasures against SAMHD1 restriction remain unknown. Using Epstein-Barr virus (EBV) as a primary model, we discovered that SAMHD1-mediated anti-viral restriction is antagonized by EBV BGLF4, a member of the conserved viral protein kinases encoded by all herpesviruses. Mechanistically, we found that BGLF4 phosphorylates SAMHD1 and thereby inhibits its dNTPase activity. We further demonstrated that the targeting of SAMHD1 for phosphorylation is a common feature shared by beta- and gamma-herpesviruses. Together, our findings uncover a unique immune evasion mechanism whereby herpesviruses exploit the phosphorylation of SAMHD1 to thwart host defenses.

**Highlights:**

- SAMHD1 depletion facilitates EBV lytic replication.
- EBV protein kinase BGLF4 directly phosphorylates SAMHD1.
- BGLF4 phosphorylation of SAMHD1 inhibits its dCTPase and dTTPase activity.
- SAMHD1 is targeted by the conserved herpesvirus protein kinases.

**In Brief:** Herpesviruses have evolved elegant strategies to dampen the host anti-viral responses. Zhang *et al.* discover a unique mechanism by which herpesviruses evade SAMHD1-mediated host defenses through phosphorylation, expanding the functional repertoire of viral protein kinases in herpesvirus biology

## Introduction

Human herpesviruses are ubiquitous pathogens that establish lifelong persistent infections and are associated with a variety of diseases ranging from cold sores to mononucleosis, birth defects, and cancers. During a long period of co-evolution with their hosts, these viruses have established multiple strategies to combat the cellular defense systems. The eight human herpesviruses discovered to date are classified into three subfamilies, alpha-, beta- and gamma-herpesviruses. The alpha-herpesviruses consist of herpes simplex 1 and 2 (HSV-1 and HSV-2) and varicella– zoster virus (VZV); the beta-herpesviruses include human cytomegalovirus (HCMV), human herpesviruses 6 (HHV-6) and human herpesvirus 7 (HHV-7); and the gamma-herpesviruses are Epstein–Barr virus (EBV) and Kaposi sarcoma-associated herpesvirus (KSHV) (Gershburg and Pagano, 2008; Li and Hayward, 2013). Although these herpesviruses express a limited number of unique latency-associated genes, they each encode a large number of functionally conserved lytic genes that are critical for efficient virus replication and spread. Among the conserved lytic gene products are the orthologous serine/threonine protein kinases, namely HSV-1/2 UL13, VZV ORF47, HCMV UL97, HHV-6/7 U69, EBV BGLF4 and KSHV ORF36 (Gershburg and Pagano, 2008; Li et al., 2011). These viral protein kinases are referred to as conserved herpesvirus protein kinases (CHPKs). CHPKs interact with and potentially phosphorylate a variety of cellular proteins (Calderwood et al., 2007; Li et al., 2015; Li et al., 2011). Although CHPKs from beta- and gamma-herpesviruses are structurally similar to the cellular cyclin dependent kinases 1/2 (CDK1 and CDK2) (Kuny et al., 2010; Romaker et al., 2006), recent studies from our lab and others suggest that these CHPKs have a broader substrate recognition than the cellular ortholog CDKs (Li et al., 2015; Li et al., 2011; Oberstein et al., 2015; Zhu et al., 2009). The mimicry of CDK activity by CHPKs results in the phosphorylation of retinoblastoma (Rb) (Hume et al., 2008; Iwahori et al., 2015), MCM4 (Kudoh et al., 2006), condensin (Lee et al., 2007), stathmin (Chen et al., 2010), EF-1 delta (Kawaguchi and Kato, 2003) and p27 (Iwahori et al., 2009), which leads to a prolonged pseudo-S-phase environment suitable for efficient viral DNA replication (Kuny et al., 2010). CHPKs actively trigger the host DNA damage response through TIP60-ATM-H2AX axis to benefit virus replication (Li et al., 2011; Tarakanova et al., 2007). CHPKs block interferon response through inhibiting IRF3 activity to facilitate viral persistence (Hwang et al., 2009; Ma et al., 2015; Wang et al., 2009). CHPKs also phosphorylate lamin A/C, nuclear pore complex and viral nuclear egress complex to enhance the egress of virus capsids from the nucleus (Chang et al., 2012; Chang et al., 2015; Hamirally et al., 2009; Lee et al., 2008; Li et al., 2015; Oberstein et al., 2015; Sharma et al., 2015; Sharma and Coen, 2014; Sharma et al., 2014). In addition to protein phosphorylation, some CHPKs are responsible for the phosphorylation of the nucleoside analog drugs acyclovir and ganciclovir in virus-infected cells (Gershburg et al., 2004; Meng et al., 2010; Moore et al., 2001; Sullivan et al., 1992). Deletion or inhibition of CHPKs activity results in the suppression of virus replication of EBV (Gershburg et al., 2007; Li et al., 2011), HCMV (Prichard et al., 1999; Wolf et al., 2001), HSV-1 (Shibaki et al., 2001) and murine gamma-herpesvirus 68 (MHV68) (Tarakanova et al., 2007). Given the important roles of CHPKs in herpesvirus infection, drugs targeting CHPKs are promising for treating herpesvirus-associated diseases. One protein kinase inhibitor, maribavir, targeting HCMV protein kinase UL97 has entered clinical trials (Prichard, 2009). Inhibitors that can decrease the expression of CHPKs also show promise in blocking virus replication (Sun et al., 2013).

Although significant progress has been made toward CHPK substrate identification, the functional importance of these CHPK substrates in virus replication remains to be determined. One potential CHPK target, sterile alpha motif and HD domain 1 (SAMHD1) was recently identified by our phospho-proteomic screening (Li et al., 2015). SAMHD1 is a newly defined anti-viral host restriction factor that limits the infection of human immunodeficiency virus-1 (HIV-1) (Laguette et al., 2011), HSV-1 (Hollenbaugh et al., 2013; Kim et al., 2013), vaccinia virus (Hollenbaugh et al., 2013), human T cell leukemia virus type 1 (Sze et al., 2013) and hepatitis B virus (HBV) (Chen et al., 2014). Phosphorylation of SAMHD1 by CDK1 and CDK2 plays a critical role in lentiviral restriction. Initial studies suggest that the anti-viral restriction function but not the dNTPase activity of SAMHD1 is controlled by phosphorylation (Kim et al., 2013; Welbourn et al., 2013; White et al., 2013). However, more recent studies demonstrate that the dNTPase activity is down-regulated but not completely inactivated by phosphorylation to finely tune the function of SAMHD1 during both viral and cellular DNA replication (Arnold et al., 2015; Badia et al., 2015; Ruiz et al., 2015; Tang et al., 2015; Wittmann et al., 2015; Yan et al., 2015). HIV-2/SIV virion-associated Vpx accessory proteins target SAMHD1 for rapid proteasome-dependent degradation to ensure efficient transduction of myeloid cells (Hrecka et al., 2011; Schwefel et al., 2015; Schwefel et al., 2014). For all other viruses restricted by SAMHD1, the viral anti-SAMHD1 restriction strategies remain to be determined.

In this study, we identify SAMHD1 as an anti-EBV restriction factor. Our results show that depletion of SAMHD1 leads to a significant increase of viral DNA replication without affecting viral protein expression. Interestingly, we demonstrate that EBV protein kinase BGLF4 interacts with and phosphorylates SAMHD1 to inhibit its dNTPase activity. More importantly, CHPKs from all three herpesvirus subfamilies interact with SAMHD1 and those from beta- and gamma-herpesviruses phosphorylate SAMHD1 in cells. Together, our results suggest that SAMHD1 is a key regulator for all herpesviruses and CHPKs are direct viral countermeasures against SAMHD1.

## Results

### SAMHD1 is phosphorylated by the EBV protein kinase BGLF4

To better understand the function of EBV protein kinase BGLF4 in the viral life cycle, we recently utilized a Stable Isotope Labeling by Amino acids in Cell culture (SILAC)-based quantitative mass spectrometry (MS) approach to identify potential targets for EBV BGLF4 (Li et al., 2015). More than 1,000 hyperphosphorylated cellular proteins were identified in BGLF4-expressing cells. Among these proteins, we detected SAMHD1. As shown in **Fig 1A**, the phosphorylation of SAMHD1 on T592 is significantly increased in BGLF4-expressing cells. To further confirm these results, we monitored the phosphorylation status of SAMHD1 in Akata (EBV+) cells carrying either control or BGLF4-expressing vectors. As expected, the phosphorylation of SAMHD1 was gradually increased with BGLF4 induction by doxycycline (**Fig 1B,** lanes 4-6). In contrast, the phosphorylation of SAMHD1 was even decreased with the same treatment in the vector control cells (**Fig 1B,** lanes 1-3). Our previous study demonstrated that EBV protein kinase BGLF4 is a Small Ubiquitin-like Modifier (SUMO) binding protein. The SUMO binding-deficient mutant BGLF4 could not trigger the DNA damage response although the BGLF4 kinase activity was not affected *in vitro* (Li et al., 2012). Here we also tested the function of this SUMO binding-deficient (m-SIM-N) mutant together with kinase-dead (KD) BGLF4 in SAMHD1 phosphorylation. Interestingly, we found that the SUMO binding-deficient (m-SIM-N) BGLF4 also triggered the phosphorylation of SAMHD1 while the expression of kinase-dead (KD) BGLF4 reduced the phospho-SAMHD1 level (**Fig 1B,** lanes 7-9 vs lanes 10-12). These results suggested that the EBV protein kinase BGLF4 triggered the phosphorylation of SAMHD1 in a kinase-activity dependent manner and the SUMO-binding function of BGLF4 was dispensable for SAMHD1 phosphorylation. By using phos-tag SDS-PAGE to separate phospho- and non-phospho-proteins (Kinoshita et al., 2006), we found that nearly all SAMHD1 is converted to the phosphorylated form in BGLF4-expressing cells (**Fig 1C**).

**Figure 1.**
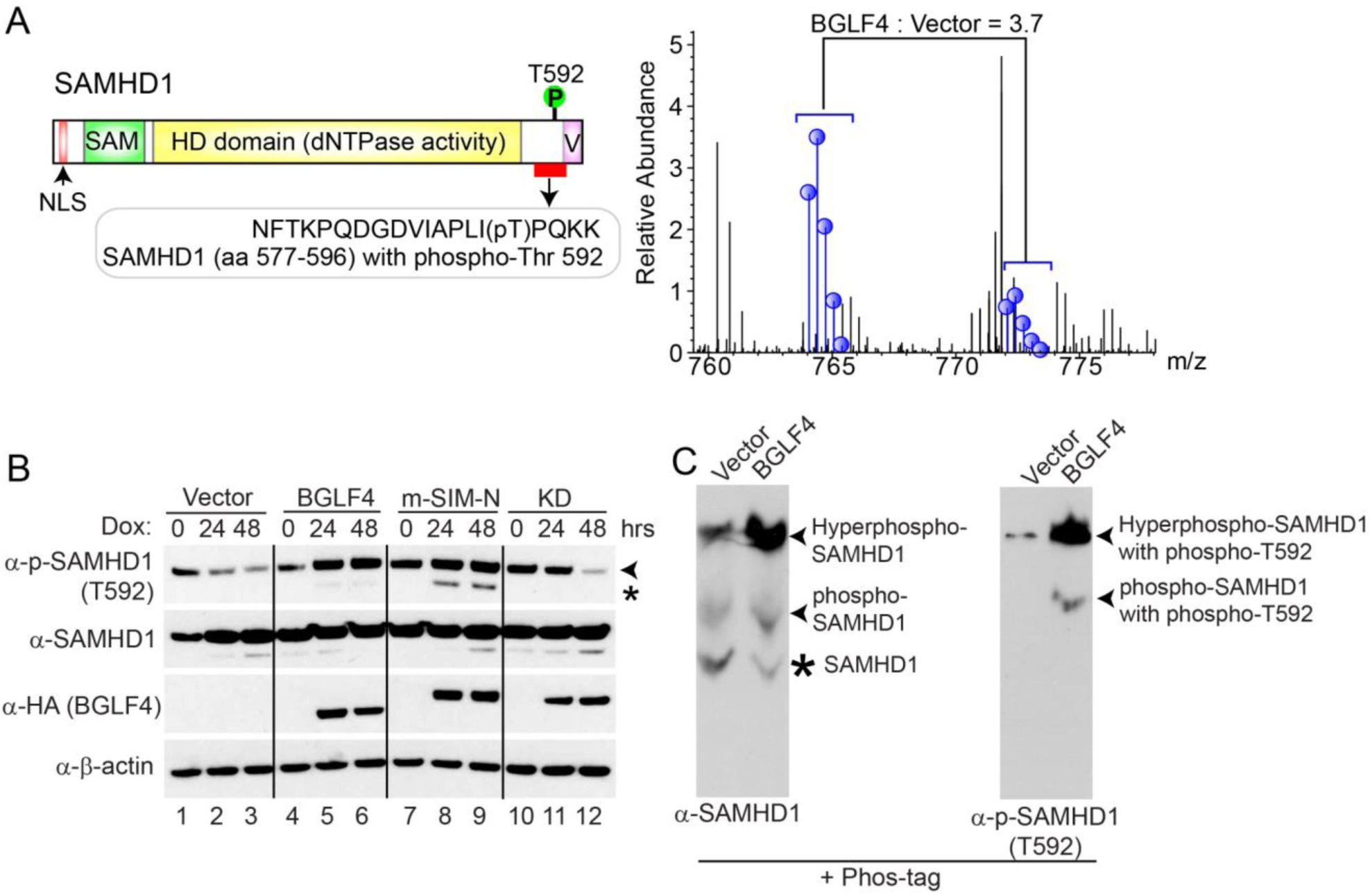
SAMHD1 regulation by EBV protein kinase BGLF4. (A) SAMHD1 is phosphorylated upon BGLF4 induction. Schematic representation of SAMHD1 protein is presented in the top panel with the relative positions for each functional doman as indicated. NLS: nuclear localization signal peptide; SAM: sterile alpha motif; HD: histidine-aspartic domain; V: Vpx interacting domain. SILAC-based quantitative proteomic analyses revealed that the SAMHD1 phosphorylation on T592 is increased upon BGLF4 induction for 48 hrs. (B) Phosphorylation of SAMHD1 is induced by wild-type and SUMO-binding-deficient mutant BGLF4. Western blot analysis was performed on cell lysates from Akata (EBV+) cells carrying control vector (Vector), wild-type, SUMO-binding-deficient (mSIM-N), or kinase-dead (KD) BGLF4 using anti-p-SAMHD1 (T592), anti-SAMHD1, anti-HA and anti-β-actin antibodies as indicated. The cells were either untreated (0 hr) or treated with doxycycline (Dox) for 24 or 48 hrs to induce BGLF4 expression. Arrow head denotes the major 72 kDa phospho-SAMHD1 band and asterisk denotes a 69 kDa non-specific band or a phospho-SAMHD1 band derived from a SAMHD1 isoform protein. (C) BGLF4 induces hyperphosphorylation of SAMHD1. Akata (EBV+) cells carrying control vector (Vector) or wild-type BGLF4 were treated with doxycycline for 48 hrs. Cell extracts were separated using Phos-tag acrylamide gel and then analyzed with Western Blot using antibodies against SAMHD1 (left) and phopho-SAMHD1 (T592) (right). Different forms of SAMHD1 and phospho-SAMHD1 are indicated.

To further test whether BGLF4-triggered phosphorylation of SAMHD1 is physiologically relevant during EBV lytic replication, we monitored the phosphorylation of SAMHD1 in lytically induced Akata-BX1 (EBV+) B cells. As a control, the same IgG crosslinking treatment was also applied to Akata-4E3 (EBV-) B cells. Consistent with the results from BGLF4-expressing cells (**Fig 1B**), we observed a time-dependent increase of phospho-SAMHD1 during the course of EBV lytic reactivation, which correlates nicely with the BGLF4 expression level (**Fig 2A,** lanes 1-4). In contrast, the phospho-SAMHD1 level gradually decreased in Akata-4E3 (EBV-) B cells following the anit-IgG treatment (**Fig 2A,** lanes 5-8). In addition to the major phospho-SAMHD1 band (72 kDa) (**Fig 2A,** arrow head), we also noticed a 69 kDa band (**Fig 2A,** asterisk). This faster migrating band was possibly derived from another isoform of SAMHD1 (Shi et al., 2014; Welbourn et al., 2012). In addition to B cells, we also monitored the phosphorylation of SAMHD1 in EBV-positive epithelial cells, Hela-(EBV+) cells (Feng et al., 2007) following transfection of EBV ZTA and RTA to induce EBV lytic replication. Interestingly, we found that the transfection of both ZTA and RTA triggers BGLF4 expression and SAMHD1 phosphorylation (**Fig 2B,** lane 3). Together, these results suggested that the phosphorylation of SAMHD1 is mediated by EBV protein kinase BGLF4.

**Figure 2.**
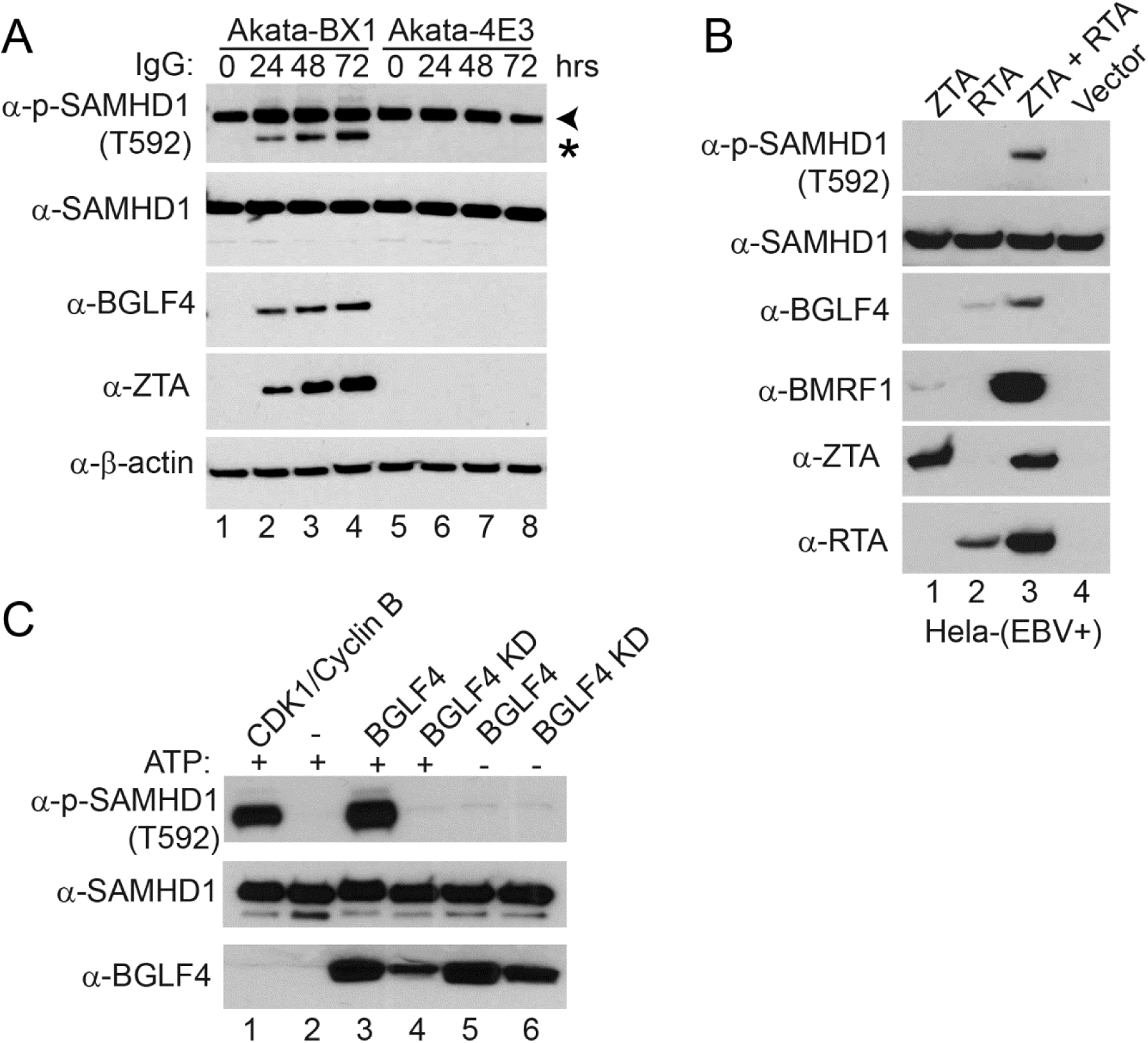
SAMHD1 is phosphorylated by EBV protein kinase BGLF4. (A) SAMHD1 is phosphorylated upon lytic induction of EBV. Western blot analysis was performed on cell lysates from Akata-BX1 (EBV+) and Akata 4E3 (EBV-) cells using anti-p-SAMHD1 (T592), anti-SAMHD1, anti-BGLF4, anti-ZTA and anti-β-actin antibodies as indicated. The cells were either untreated (0 hr) or treated with anti-human IgG for 24, 48 or 72 hrs to induce lytic reactivation. Arrow head denotes the major 72 kDa phospho-SAMHD1 band and asterisk denotes a 69 kDa non-specific band or a phospho-SAMHD1 band derived from a SAMHD1 isoform protein. (B) SAMHD1 is phosphorylated in EBV-replicating Hela cells. Western blot analysis was performed on cell lysates from Hela (EBV+) transfected with EBV ZTA, RTA or ZTA plus RTA as indicated for 48 hrs to induce lytic reactivation. The phosphorylation of SAMHD1 correlated with BGLF4 expression. (C) SAMHD1 is phosphorylated by EBV protein kinase BGLF4 *in vitro*. Recombinant SAMHD1 protein was mixed with purified wild-type or kinase-dead (KD) BGLF4 for 30 min at 30 °C. As a positive control, SAMHD1 was incubated with CDK1/Cyclin B (lane 1), which is a known kinase for SAMHD1. As negative controls, either kinase or ATP was omitted in the reaction mixture (lanes 2, 5, 6). The phospo-SAMHD1, SAMHD1 and BGLF4 were detected by Western Blot using antibodies as indicated.

To unambiguously prove SAMHD1 as a direct substrate for EBV protein kinase BGLF4, we performed an *in vitro* kinase assay. We found that the wild-type EBV protein kinase BGLF4, but not the kinase-dead (KD) mutant, phosphorylates on SAMHD1 T592, which is similar to the phosphorylation induced by the kinase complex CDK1/Cyclin B (Welbourn et al., 2013) (**Fig 2C,** lanes 1, 3 versus 4). In contrast, we didn’t detect the phosphorylation signal in reactions lacking BGLF4 or ATP (**Fig 2B,** lanes 2, 5 and 6). Taken together, our data suggested that EBV protein kinase BGLF4 phosphorylates SAMHD1 *in vivo* and *in vitro*.

### SAMHD1 restricts EBV replication

Because SAMHD1 plays a critical role in restricting HIV and HSV replication and the phosphorylation on T592 can block it anti-viral restriction function, we hypothesized that SAMHD1 is also a restriction factor for EBV. To test our hypothesis, we examined whether SAMHD1 expression affected viral protein expression. Akata-BX1 (EBV+) cells were infected with individual sg-RNA lentivirus to deplete SAMHD1 by CRISPR/Cas9-meidated genome editing. The depletion efficiency of SAMHD1 was high for both sg-RNAs compared with control (**Fig 3B,** lanes 1-6 vs lanes 7-9). We measured accumulation of the immediate-early and early viral proteins ZTA and BGLF4 during the course of EBV lytic reactivation. The levels of viral proteins in SAMHD1-depleted cells were similar to those seen in control cells, suggesting that SAMHD1 did not affect the expression of viral proteins (**Fig 3B,** ZTA and BGLF4 blots). To test whether SAMHD1 plays a role in EBV replication, we measured the infectious EBV particles following lytic induction by IgG crosslinking. Interestingly, we found that the relative EBV titer were significantly higher in SAMHD1-knockout cells than those in the control cells at 48 hrs post-lytic induction (**Fig 3B,** lower panel, lanes 3 & 6 verus 9). In the EBV-replicating Akata cells, the cellular CDK1 protein level was dramatically reduced upon anti-IgG treatment (**Fig 3B,** CDK1 blot), suggesting that SAMHD1 phosphorylation was not mediated by CDK1 but mainly mediated by the EBV protein kinase BGLF4 (**Fig 3B,** phospho-SAMHD1 and BGLF4 blots, lanes 7-9).

**Figure 3.**
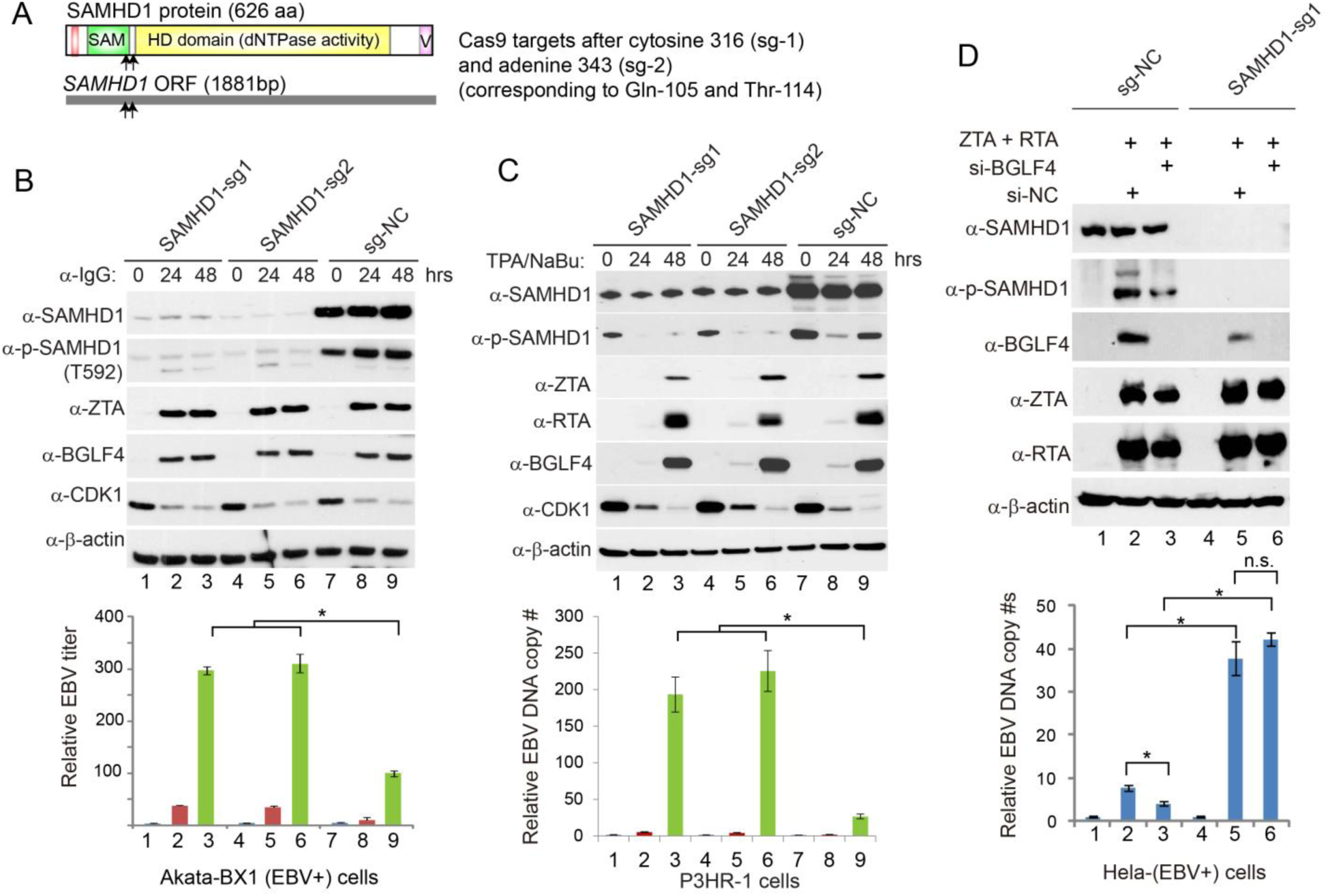
SAMHD1 depletion facilitates EBV lytic replication. (A) Schematic representation of Cas9 target sites within the 1881 bp *SAMHD1* open read frame (ORF). The corresponding positions of amino acids are also shown. (B) SAMHD1 depletion facilitates EBV replication in Akata cells. SAMHD1-depleted (sg-1 and sg-2) and control (sg-NC) Akata-BX1 (EBV+) B cells were treated with IgG cross-linking to induce EBV lytic reactivation. SAMHD1 protein and phosphorylation levels, the accumulation of EBV ZTA and BGLF4, and the CDK1 protein level were analyzed by immunoblotting at the indicated time points. β-actin served as a loading control. Relative EBV titer of lytically induced Akata-BX1 (EBV+) cells carrying SAMHD1-deletion or control was measured using a Raji cell infection assay. (C) SAMHD1 depletion facilitates EBV replication in P3HR-1 cells. SAMHD1-depleted (sg-1 and sg-2) and control (sg-NC) P3HR-1cells were treated with TPA and Sodium Butyrate (NaBu) to induce EBV lytic reactivation. Western blots were performed using antibodies as indicated. β-actin served as a loading control. Relative EBV DNA copy numbers were determined by qPCR using primers to *BALF5* gene normalized by *β-actin*. (D) BGLF4 knockdown suppresses EBV replication in SAMHD1-expressing cells but not in SAMHD1-depleted cells. Control (sg-NC) and SAMHD1-depleted (sg-1) Hela (EBV+) cells were transfected with ZTA plus RTA to induce EBV lytic reactivation. The siBGLF4-expressing plasmid was co-transfected to knockdown BGLF4. The transfection of si-NC served as a negative control for siBGLF4. The EBV genome copy numbers were measured by qPCR using primers specific to EBV *BALF5* gene normalized by *β-actin*. Representative results from three biological replicates are presented. Error bars indicate the standard deviation. * p < 0.05; n.s.: not significant. See also Figure S1.

To further demonstrate SAMHD1-mediated restriction toward EBV replication, we depleted SAMHD1 by a similar CRISPR/Cas9 method in the parental Akata (EBV+) and the P3HR-1 cells. Similarly, knocking down SAMHD1 promoted EBV DNA replication in the parental Akata (EBV+) cells (**Fig S1**). In P3HR-1 cells, we also observed that SAMHD1 depletion significantly enhanced EBV DNA replication with minimal effects on viral protein expression following TPA and sodium butyrate treatment (**Fig 3C,** lanes 3 & 6 versus 9), reinforcing that SAMHD1 plays a key role in restricting EBV replication. Similar as SAMHD1 de-phosphorylation observed in THP-1 cells differentiated by TPA treatment (Cribier et al., 2013; Kim et al., 2013; Welbourn et al., 2013), we also noticed an initial SAMHD1 de-phosphorylation upon TPA and sodium butyrate treatment of P3HR-1 cells (**Fig 3C,** phospho-SAMHD1 blot, lane 7 versus 8). However, SAMHD1 phosphorylation re-emerged at 48 hrs post-treatment when EBV protein kinase BGLF4 was highly expressed while cellular CDK1 level was significantly reduced (**Fig 3C,** phospho-SAMHD1 blot, lane 9), further suggesting BGLF4 may be responsible for the re-occurrence of SAMHD1 phosphorylation.

To further evaluate the function of SAMHD1 and its regulation by EBV BGLF4 in EBV lytic replication, we created a SAMHD1 depletion cell line using Hela-(EBV+) cells. The advantage of using Hela-(EBV+) cells is that EBV lytic replication can be triggered by ZTA and RTA transfection and that BGFL4 can be easily knocked down by co-transfection of a plasmid encoding siBGLF4 (Gershburg et al., 2007) to evaluate the role of BGLF4-SAMHD1 axis in EBV replication. As expected, ZTA/RTA transfection triggered EBV lytic replication and SAMHD1 phosphorylation while BGLF4 knocking down led to reduced SAMHD1 phosphorylation and EBV replication (**Fig 3D,** lanes 2 and 3). SAMHD1 depletion significantly enhanced EBV replication and, interestingly, BGLF4 knocking down did not affect viral replication in SAMHD1-depleted cells **(Fig 3D,** lanes 2 and 3), suggesting that BGLF4 promotes EBV DNA replication mainly through inhibiting SAMHD1.

### Phospho-mimicking SAMHD1 promotes while phospho-deficient SAMHD1 blocks EBV lytic replication

To further prove BGLF4-mediated SAMHD1 phosphorylation is important for EBV lytic replication, we created a phospho-mimicking (T592D) and a phospho-deficient (T592A) SAMHD1. We then used the SAMHD1-depleted Akaka (EBV+) cells to reconstitute wild-type and the mutant SAMHD1. During the course of EBV lytic reactivation, the levels of viral proteins were similar among these cell lines, suggesting that SAMHD1 and its phosphorylation status did not affect the expression of viral proteins (**Fig 4,** upper panel). As expected the reconstitution of wild-type SAMHD1 suppressed EBV DNA replication (**Fig 4,** lower panel, lane 3 versus 6). Interestingly, the reconstitution of phospho-mimicking SAMHD1 (T592D) promoted EBV DNA replication compared with cells carrying the vector control or WT SAMHD1 (**Fig 4,** lower panel, lane 9 versus 3 and 6). The higher DNA replication in phospho-mimicking (T592D) cells compared with the SAMHD1-depleted cells suggested that phosphorylation of SAMHD1 on T592 may even promote DNA repair and/or fork progression to promote viral DNA replication (Coquel et al., 2018; Daddacha et al., 2017). In contrast, the phospho-deficient SAMHD1 (T592A) blocks EBV DNA replication compared with cells carrying WT SAMHD1 or T592D mutant (**Fig 4,** lower panel, lane 12 versus 6 and 9). Together, these results indicated that SAMHD1 phosphorylation by a viral kinase plays a critical role in promoting EBV DNA replication.

**Figure 4.**
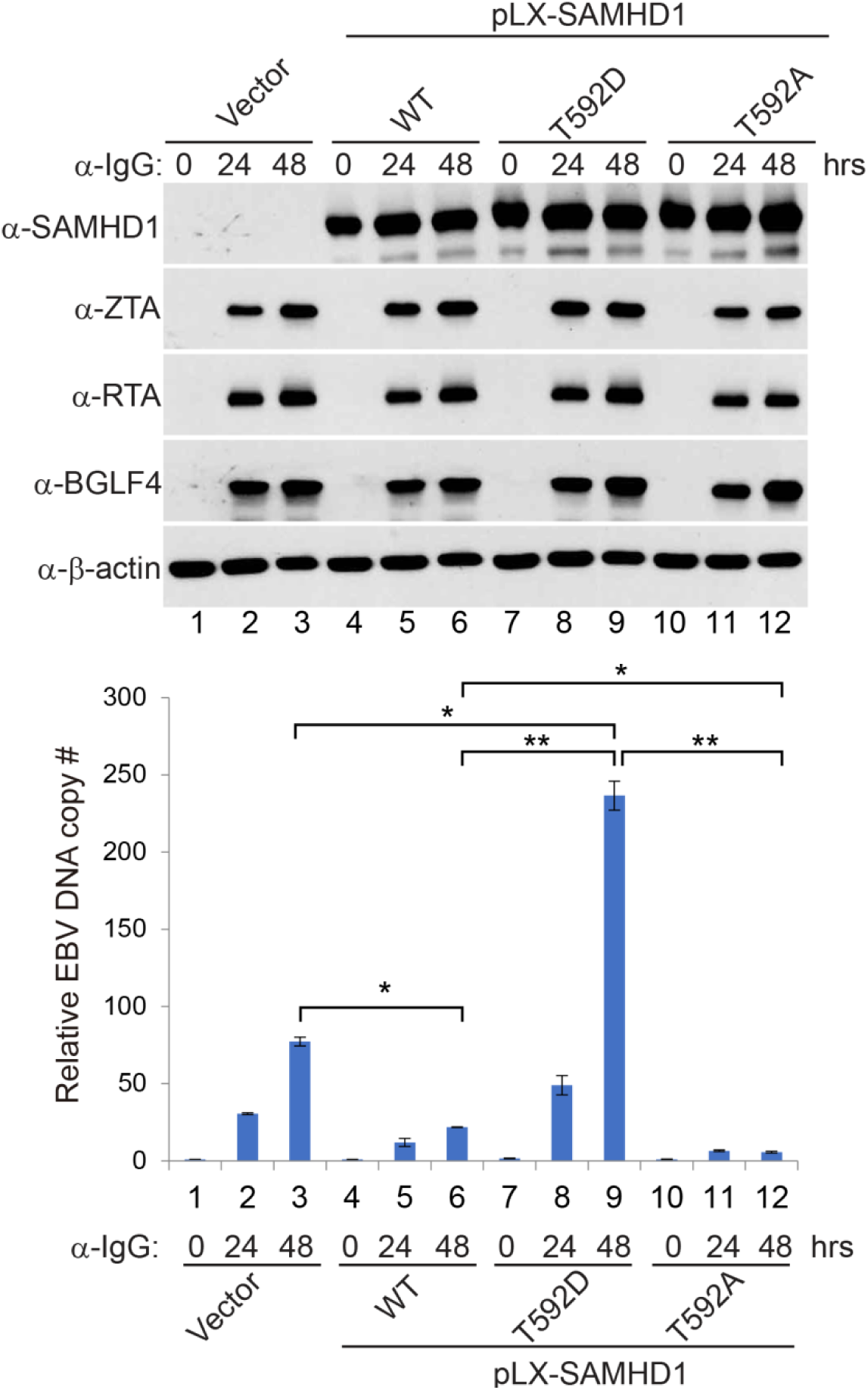
SAMHD1 reconstitution suppresses EBV lytic replication. SAMHD1-depleted (sg1) Akata (EBV+) cells were used to establish SAMHD1-expressing stable cell lines using pLX-304 lentiviral constructs containing wild-type (WT), T592D and T592A SAMHD1. Western blot analyses showing SAMHD1, ZTA, RTA and BGLF4 expression level in different cell lines upon IgG cross-linking as indicated. Viral DNA replication was measured by qPCR using primers to *BALF5*. Representative results from three biological replicates are presented. The value of Vector control at 0 hr (lane 1) was set as 1. Error bars indicate the standard deviation. *p < 0.05; ** p < 0.001.

### EBV BGLF4 inhibits SAMHD1 dNTPase activity through phosphorylation

We hypothesized that EBV protein kinase BGLF4 can alleviate the anti-viral activity of SAMHD1 through phosphorylation. To get mechanistic insights into how BGLF4 regulates SAMHD1, we measured the dNTPase activity of SAMHD1 and phospho-SAMHD1 triggered by BGLF4. SAMHD1 was first incubated with BGLF4, kinase dead BGLF4 (BGLF4-KD), CDK1/CyclinB or buffer control and then SAMHD1 and phospho-SAMHD1 were activated by incubating with GTP and a mixture of all dNTP at equal concentration. Subsequently, each individual dNTP was added to the activated SAMHD1 mixture for testing SAMHD1’s dNTPase activity. Interestingly, we found that phospho-SAMHD1 triggered by BGLF4 and CDK1/Cyclin B shows reduced dCTPase and dTTPase activity while the dATPase and dGTPase activity was less affected **(Fig 5**). The results suggested that EBV BGLF4 promotes viral replication through selectively inhibiting SAMHD1’s dNTPase activity.

**Figure 5.**
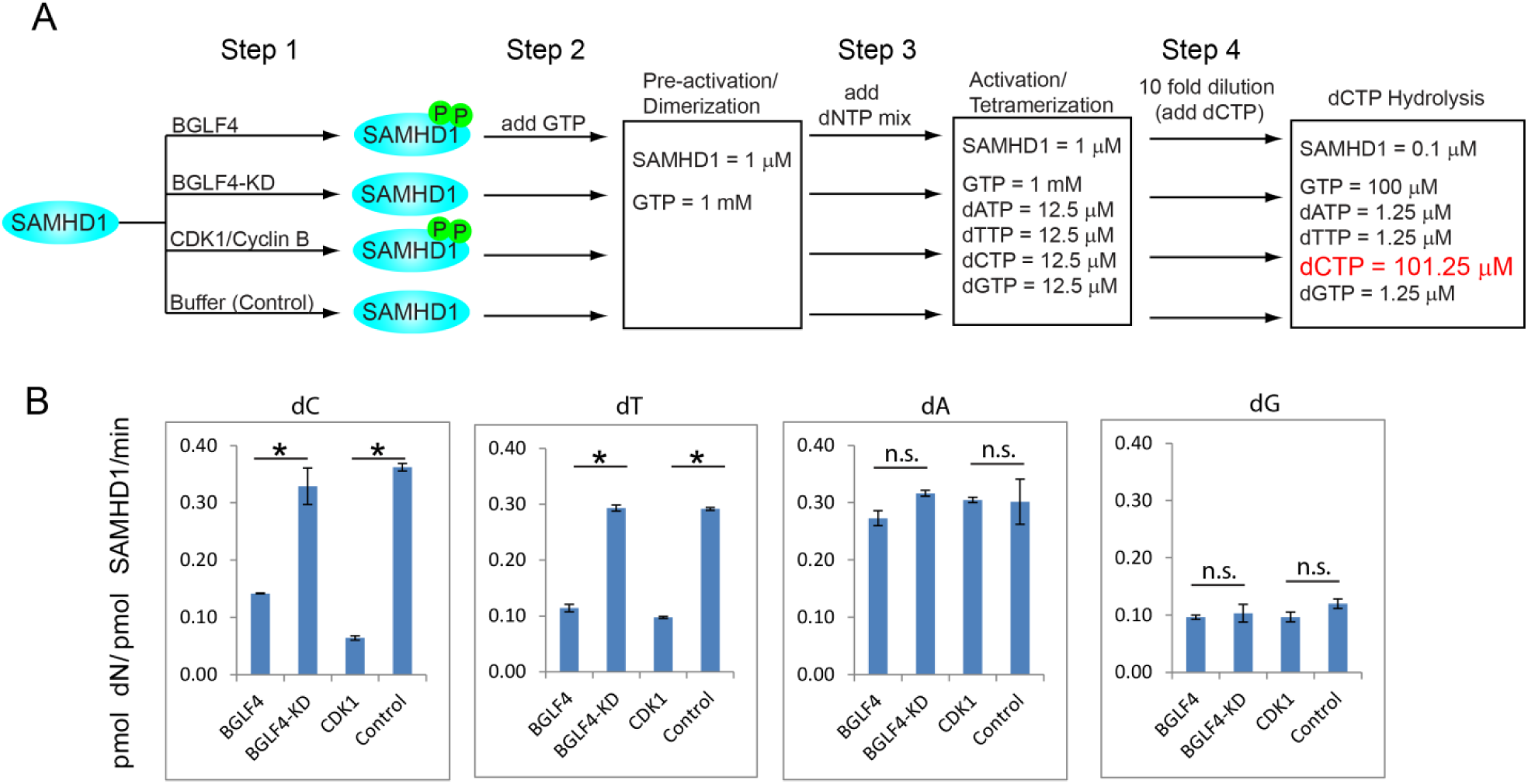
BGLF4 suppresses the dCTPase and dTTPase activity of SAMHD1 through phosphorylation. (A) Schematic representation of SAMHD1 phosphorylation, activation and dCTP hydrolysis analysis. Step 1: Recombinant SAMHD1 was phosphorylated by BGLF4 or CDK1 *in vitro*. Non-phosphorylated SAMHD1 was similarly processed by incubating with kinase-dead BGLF4 (BGLF4-KD) or Buffer control. Step 2: The phosphorylated and non-phosphorylated SAMHD1 were converted to GTP-bound dimer. Step 3: SAMHD1 dimer was further activated by adding dNTP mix for 60 sec. Step 4: The mixture was diluted 10-fold and the substrate dCTP is added for 0 to 240 min. For analysis of other dNTP hydrolysis, 100 μM dTTP, dATP or dGTP was added in Step 4. (B) The SAMHD1 dNTPase activity (dN generation rate, pmol dN/pmol-SAMHD1/min) was analyzed by HPLC using a Synergi C18 column 150 x 4.6 mm. Error bars represent the standard error of the mean from two independent experiments. * p<0.05; n.s.: not significant.

### Common targeting of SAMHD1 by the conserved herpesvirus protein kinases (CHPKs)

The eight human herpesviruses diverged around 100 to 200 million years ago, but they each encode a CHPK (McGeoch et al., 1995). Because the kinase domain of EBV BGLF4 shares high sequence homology with other CHPKs (**Fig 6A**), we reasoned that the targeting of SAMHD1 might be a common feature for all CHPKs. To test this possibility, we transfected 7 CHPKs representative of the alpha-, beta- and gamma-herpesvirus subfamilies into 293T cells and examined the phosphorylation status of endogenous SAMHD1 in individual CHPK-transfected cells. Interestingly, we found that wild-type CHPKs from all beta- and gamma-herpesviruses (HHV-6/7 U69, HCMV UL97, EBV BGLF4, and KSHV ORF36) induced a strong phosphorylation of SAMHD1 compared to their corresponding kinase-dead (KD) mutants and non-transfection controls (**Fig 6B**, p-SAMHD1 (T592) blot, lanes 3, 5, 7, 9 and 11 vs lanes 4, 6, 8, 10, 12 and 13). In contrast, the alpha-herpesvirus CHPKs (HSV-1 UL13 and VZV ORF47) were unable to induce SAMHD1 phosphorylation on T592 (**Fig 6B**, lanes 1 and 2). The total SAMHD1 protein level was not affected by all CHPKs, suggesting that the CHPKs induced increase of SAMHD1 phosphorylation was independent of its steady-state level. We didn’t notice a correlation between CHPKs expression level and their ability to phosphorylate SAMHD1. For example, CHPKs with high expression levels either did (HHV-6 U69, HCMV UL97, EBV BGLF4 and KSHV ORF36) or did not (VZV ORF47) trigger the phosphorylation of SAMHD1. CHPKs with low expression levels either did (HHV-7 U69) or did not (HSV-1 UL13) trigger the phosphorylation of SAMHD1 (**Fig 6B**). These results suggested that beta- and gamma-herpesvirus CHPKs, but not the alpha-herpesvirus CHPKs, phosphorylate the T592 residue of SAMHD1.

**Figure 6.**
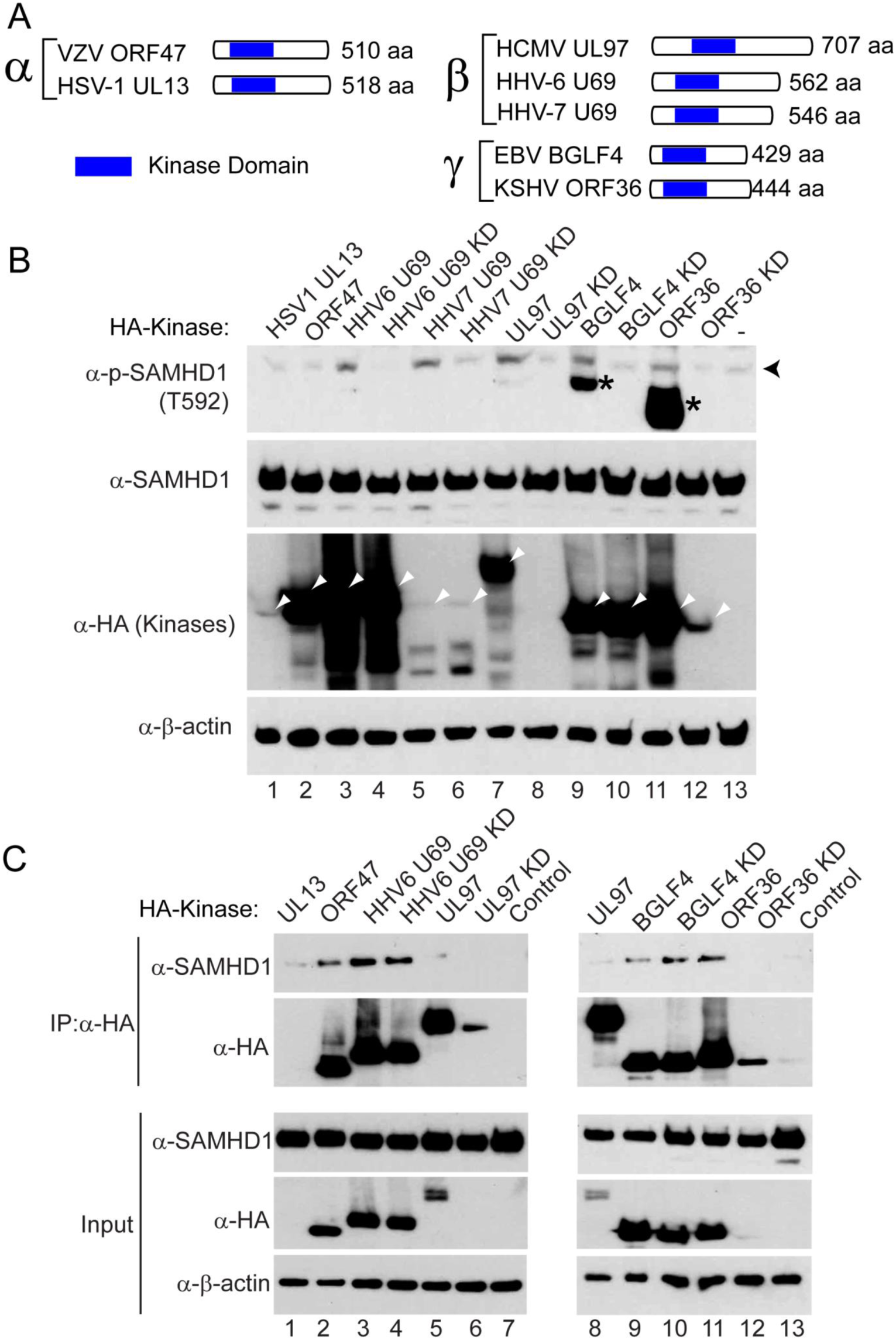
Conserved herpesvirus protein kinases target SAMHD1. (A) Schematic representation of the conserved herpesvirus protein kinases (CHPKs) from alpha-, beta- and gamma-herpesviruses. The relative position of the conserved kinase domain was shown. (B) Beta- and gamma-herpesvirus CHPKs induce the phosphorylation of SAMHD1. Western blot analysis of cell extracts showing that the phosphorylation of SAMHD1 on T592 was induced by overexpression of wild-type HHV-6/7 U69, HCMV UL97, EBV BGLF4, and KSHV ORF36. The 293T cells were transfected with individual CHPKs and kinase-dead (KD) mutants and the cells were harvested 48 hrs post-transfection. Black arrow head denotes the major 72 kDa phospho-SAMHD1 band and asterisks denote non-specific bands or phospho-SAMHD1 bands derived from a SAMHD1 isoform protein. The positions of CHPKs were indicated by white arrow heads. β-actin served as a loading control. (C) Alpha-, beta- and gamma-herpesvirus CHPKs interact with SAMHD1. Western blot analysis showing Co-IP of SAMHD1 with alpha-, beta- and gamma-herpesvirus CHPKs using 293T cells transfected with SAMHD1 and CHPKs. Input, 2% whole-cell lysate used for IP.

To test whether CHPKs interact with SAMHD1 in cells, we transfected CHPKs into 293T cells and then performed co-immunoprecipitation experiments. We found that VZV ORF47, HHV6 UL69, EBV BGLF4 and KSHV ORF36 all interacted with SAMHD1 (**Fig 6C**, lanes 2, 3, 5, 8, 9 and 11). Kinase-dead (KD) HHV-6 U69 and EBV BGLF4 preserved the interactions with SAMHD1 (**Fig 6C**, lanes 4 and 10), suggesting that the kinase activity was not required for the interaction. We detected weak or no interactions for SAMHD1 with HSV1 UL13, HCMV UL97/UL97 KD mutant and KSHV ORF36 KD mutant, possibly due to their low expression levels compared to other viral kinases (**Fig 6C**, lanes 6 and 12).

To further test whether other CHPKs directly phosphorylate SAMHD1, we performed *in vitro* kinase assays using purified CHPKs from HCMV and HSV-1. We found that the wild-type HCMV protein kinase UL97, but not the kinase-dead (KD) mutant, triggered a strong phosphorylation of SAMHD1 on T592, similar to that induced by EBV BGLF4 and CDK1/Cyclin B (**Fig 7A**, lanes 1, 2 and 8). In contrast, we didn’t detect the phosphorylation signal in reactions containing wild-type HSV-1 UL13 *in vitro* (**Fig 7A**, lane 4). To test whether UL97-induced phosphorylation of SAMHD1 is physiologically relevant during HCMV infection, we monitored the phosphorylation of SAMHD1 in HCMV infected cells. As show in **Fig 7B**, HCMV infection also triggered the phosphorylation of SAMHD1 on T592. In addition, we also confirm the previous observation that HCMV infection lead to an increase in CDK1 protein level (Gill et al., 2012). These results suggest that HCMV infection may trigger the phosphorylation of SAMHD1 by both the viral protein kinase UL97 and the cellular CDK1.

**Figure 7.**
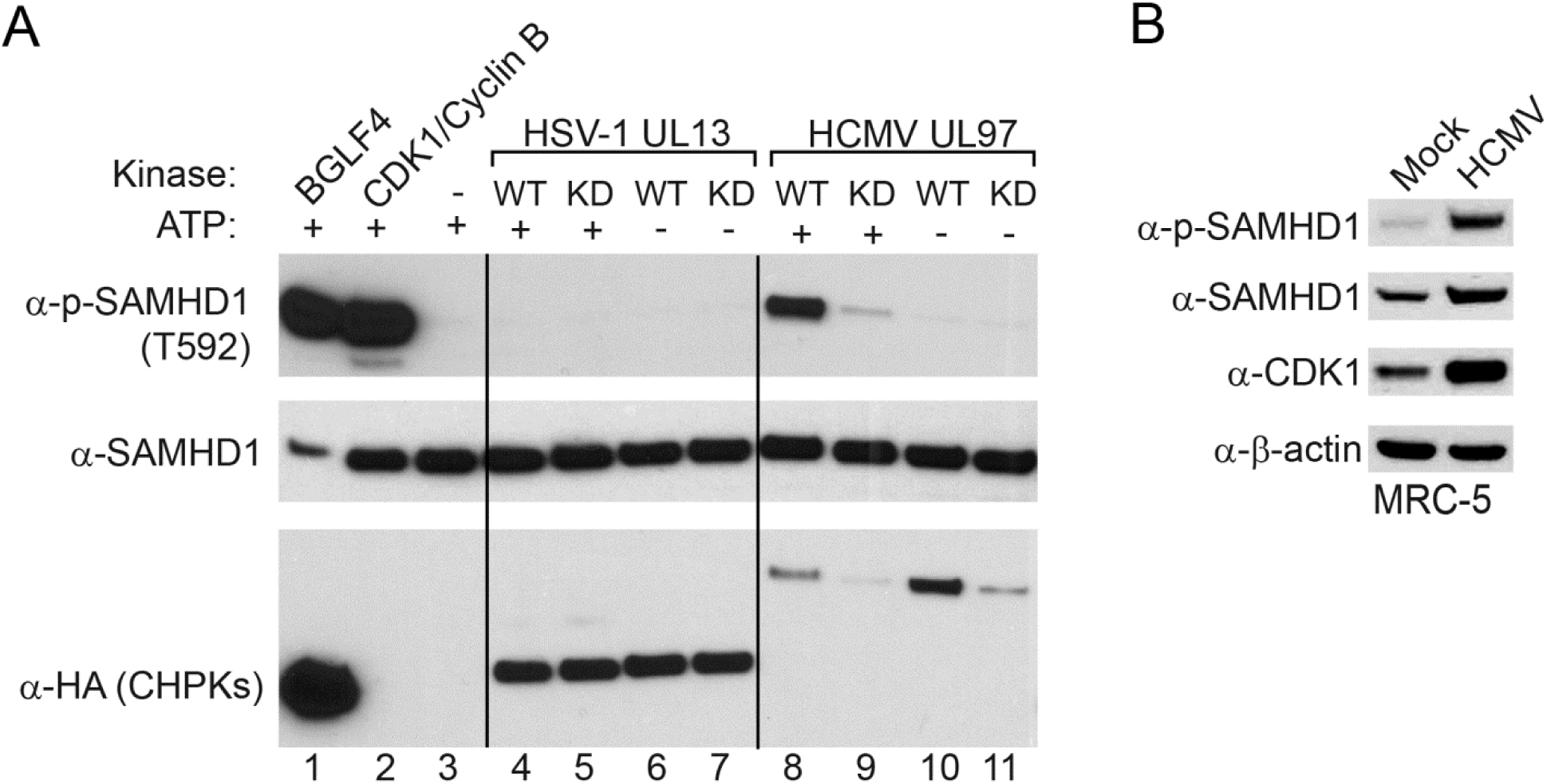
SAMHD1 is phosphorylated by HCMV protein kinase *in vitro* and upon HCMV infection in cells. (A) SAMHD1 is phosphorylated by HCMV protein kinase UL97. Recombinant SAMHD1 protein was mixed with purified wild-type (WT) or kinase-dead (KD) CHPKs as indicated for 30 min at 30 °C. As positive controls, SAMHD1 was incubated with BGLF4 or CDK1/Cyclin B. As negative controls, either CHPKs or ATP were omitted in the reaction mixture. The phospo-SAMHD1, SAMHD1 and CHPKs were detected by Western Blot using antibodies as indicated. (B) SAMHD1 is phosphorylated upon HCMV infection. Western blot analysis was performed on cell lysates from MRC-5 cells using antibodies as indicated. The cells were either mock infected or infected by HCMV (MOI=1) for 72 hrs.

Taken together, our results suggested that antagonizing SAMHD1-mediated anti-viral activity by phosphorylation is evolutionarily conserved for beta- and gamma-herpesviruses.

## Discussion

The alpha-, beta- and gamma-herpesviruses have co-evolved with human over long periods of time (McGeoch et al., 1995). Despite the numerous cellular defense mechanisms to prevent viral replication, herpesvirues are able to replicate efficiently. Not only have the viruses evolved ways to dampen the immune responses, but also have elegant strategies to counteract host restriction factors.

SAMHD1, originally identified as a human homologue of mouse interferon-γ induced protein (Li et al., 2000), is a restriction factor for a variety of viruses including HIV-1, HSV-1, vaccinia virus, and HBV (Chen et al., 2014; Hollenbaugh et al., 2013; Kim et al., 2013; Laguette et al., 2011). SAMHD1 also plays important roles in DNA repair and the degradation of nascent DNA at stalled replication forks (Coquel et al., 2018; Daddacha et al., 2017). We have shown that SAMHD1 depletion leads to a significant increase in viral replication for EBV (**Fig 3 and Fig S1**). We have demonstrated that most CHPKs counteract the restriction of SAMHD1 by phosphorylation, indicating that beta- and gamma-herpesviruses have evolved within their conserved kinase an anti-SAMHD1 mechanism to foster viral DNA replication.

In contrast to HIV-2 and the related simian immunodeficiency viruses (SIVs), which encode Vpx proteins to antagonize SAMHD1 through ubiquitin-proteasome dependent degradation (Hrecka et al., 2011; Lenzi et al., 2015), the beta- and gamma-herpesviruses utilize CHPKs to phosphorylate SAMHD1 on T592 without affecting its protein level (**Figs 1, 2 and 6**). The phosphorylation of SAMHD1 on T592 plays an important role in blocking the restriction of SAMHD1 toward HIV-1 (Arnold et al., 2015; Cribier et al., 2013; Tang et al., 2015; White et al., 2013; Yan et al., 2015). In a transfection-based overexpression system, it appears that the dNTPase activity is not regulated by phospho-mimicking SAMHD1 mutants (Kim et al., 2013; White et al., 2013). However, more recent studies supports that the phosphorylation of SAMHD1 on T592 could lead to the inhibition of its dNTPase activity in low dNTP level conditions (Arnold et al., 2015; Badia et al., 2015; Ruiz et al., 2015; Tang et al., 2015; Yan et al., 2015). Similar as cellular CDK1/Cyclin B (Jang et al., 2016), we show evidence that EBV protein kinase BGLF4 phosphorylates SAMHD1 and inhibits its dCTPase and dTTPase activity *in vitro* (**Fig 5**). During herpesvirus lytic reactivation, the massive viral DNA replication may require a large amount of dNTPs. Therefore, the CHPKs induced phosphorylation of SAMHD1 would ensure the dNTPs supply for viral DNA replication.

Because T592 of SAMHD1 is the target site of cellular CDK1 and CDK2, our results further support the previous observation that CDK-like activity is shared by beta- and gamma-herpesvirus CHPKs (Kuny et al., 2010). Although only CHPKs form beta- and gamma-herpesviruses triggered the phosphorylation of SAMHD1 on T592, the interaction of alpha-herpesvirus CHPKs with SAMHD1 suggested that these kinases may also regulate SAMHD1 activity without affecting T592 phosphorylation level or with phosphorylation on different sites (**Figs 6 and S2**). Alternatively, the alpha-herpsvirus CHPKs may coordinate with a second protein kinase (US3 from HSV1/2 and ORF66 from VZV) (Erazo and Kinchington, 2010; Kato et al., 2006) within this subfamily to induce the phosphorylation of SAMHD1.

In summary, our data suggest that SAMHD1 is a restriction factor for EBV and SAMHD1 is antagonized by CHPKs. Our findings about the mechanistic linkage between CHPKs and SAMHD1 have wide-ranging implications, especially in the discovery of new viral countermeasures for SAMHD1. Furthermore, the broad anti-viral activity of SAMHD1 toward the infection of HIV, vaccinia virus, HBV and herpesviruses suggests that SAMHD1 might control the replication of other viruses that require dNTPs to complete their life cycles and these viruses may also have evolved with similar or different anti-SAMHD1 mechanisms.

## EXPERIMENTAL PROCEDURES

### Cell Culture, Reagents and Constructs

The Akata (EBV+)-tet-Vector, Akata (EBV+)-tet-BGLF4, Akata (EBV+)-tet-BGLF4 (mSIM-N), Akata (EBV+)-tet-BGLF4 (KD), Akata (EBV+), Akata-4E3 (EBV-) cells (gifts from Diane Hayward), Akata-BX1 (EBV+) (a gift from Lindsey Hutt-Fletcher) and P3HR-1 (CAT# HTB-62, ATCC) cells were grown in RPMI 1640 media supplemented with 10% FBS (Cat# 26140079, Thermo Fisher Scientific) in 5% CO_2_ at 37 °C (Li et al., 2015; Li et al., 2012; Li et al., 2011; Lv et al., 2018; Zhang et al., 2017). 293T cells and Hela-(EBV+) cells (a gift from Shannon Kenney, University of Wisconsin-Madison) (Feng et al., 2007) were grown in DMEM media supplemented with 10% FBS (Cat# 26140079, Thermo Fisher Scientific) in 5% CO_2_ at 37 °C. P3HR-1 cells were purchased from ATCC (CAT# HTB-62). Human MRC-5 fibroblasts (ATCC CCL-171) were obtained from ATCC and propagated in high-glucose Dulbecco’s modified Eagle’s medium (Gibco BRL) supplemented with 10% fetal calf serum, 10,000 IU/liter penicillin, and 10 mg/liter streptomycin (Gibco-BRL).

The construction of Akata (EBV+)-tet-Vector, Akata (EBV+)-tet-BGLF4, Akata (EBV+)-tet-BGLF4 (mSIM-N), and Akata (EBV+)-tet-BGLF4 (KD) cell lines was described previously (Li et al., 2012; Zhu et al., 2009). The HA-tagged viral kinases (CHPKs) were gifts from Robert Kalejta (Addgene plasmid #s 26689-26698) (Kuny et al., 2010). SAMHD1 clone (Cat# SC114650) and recombinant SAMHD1 protein (Cat# TP306013) were purchased from Origene. Halo-BGLF4 (WT), Halo-BGLF4 (KD), Halo-UL97 (WT), Halo-UL97 (KD), Halo-UL13 (WT) and Halo-UL13 (KD) were constructed by cloning WT and KD kinases into pHTN HaloTag CMV-neo Vector (Cat# G7721, Promega) (Li et al., 2015). The full-length SAMHD1 was cloned into a pGEX-5x-2 vector (pGEX-5x-2-SAMHD1) with an N-terminal GST tag and Factor Xa cleavage site using Gibson assembly methods as we previous described (Zhang et al., 2017). pHTPsiRNA-PK (siBGLF4) and control pHTPsiRNA-NC (siNC) plasmids were kindly provided by Edward Gershburg (Southern Illinois University) (Gershburg et al., 2007).

### SAMHD1 Depletion by CRISPR/Cas9 Genome Editing

To knock out SAMHD1, two different sgRNAs targeting human SAMHD1 were designed and cloned into lentiCRISPR v2 vector (a gift from Feng Zhang; Addgene plasmid # 52961) (Sanjana et al., 2014). Packaging 293T cells were transfected with SAMHD1 sgRNAs or negative controls (non-targeting sgRNA-NC and sgRNA-NC2) and helper vectors (pMD2.G and psPAX2; gifts from Didier Trono; Addgene plasmid #s 12259 and 12260) using Lipofectamine 2000 reagent (Cat# 11668019, Life Technologies). Medium containing lentiviral particles and 8 mg/mL polybrene (Sigma-Aldrich, St. Louis) was used to infect Akata (EBV+) cells, Akata-BX1 (EBV+) cells, Hela-(EBV+) cells and P3HR-1 cells. Infected cells were selected in medium containing 2 μg/mL puromycin. The target guides sequences are as follows:

SAMHD1-sg1; F: 5’-caccgcttagttatatccagcgat-3’; R: 5’-aaacatcgctggatataactaagc-3’;

SAMHD1-sg2; F: 5’-caccgaatccacgttgatacaatga-3’; R: 5’-aaactcattgtatcaacgtggattc-3’;

sgRNA-NC; F: 5’-caccgttcctaagatttttaagact-3’; R: 5’-aaacagtcttaaaaatcttaggaac-3’.

### Lentiviral Transduction of SAMHD1

The pLX304-SAMHD1 was purchased from DNASU Plasmid Repository. The CRISPR-resistant SAMHD1 (nucleotide G321A, silent mutation) was generated by site-directed mutagenesis using the QuikChange II site-Directed Mutagenesis Kit (Stratagene) according to the manufacturer’s instructions. Subsequently, the pLX304-SAMHD1 T592D and T592A mutants were generated individually using the CRISPR-resistant SAMHD1 as a template. Primers sequences are: 5’-tccagcgattagttcaaatccacgttgatacaatgaag-3’ (forward) and 5’-tcaacgtggatttgaactaatcgctggatataactaag-3’ (reverse) for SAMHD1 (G321A) mutant; 5’-atgttatagccccactcatagatcctcaaaaaaaggaatggaacgacag-3’ (forward) and 5’-tcctttttttgaggatctatgagtggggctataacatcgccatcctgcg-3’ (reverse) for SAMHD1 (T592D) mutant; 5’-tgttatagccccactcatagcacctcaaaaaaaggaatg-3’ (forward) and 5’-tttgaggtgctatgagtggggctataacatcgccatcctg-3’ (reverse) for SAMHD1 (T592A) mutant. To prepare lentiviruses, 293T cells were transfected with lentiviral vector pLX304 containing the gene of wild-type SAMHD1, T592D or T592A mutants and the help vectors (pMD2.G and psPAX2) using Lipofectamine 2000 reagent. The supernatants were collected 48 hrs after transfection and used for infection of SAMHD1-depleted (sg1) cells. Infected cells were selected in medium containing 10 µg/mL blasticidin. Expression of SAMHD1 was examined by western blot analysis.

### Cell Lysis and Immunoblotting

Cells were harvested and lysed in 2x SDS-PAGE sample buffer and boiled for 5 minutes. The samples were separated on 4-20% TGX gels (Cat# 4561096, Biorad), transferred onto PVDF membranes, and probed with primary and horseradish peroxidase-conjugated secondary antibodies. Primary antibodies purchased from Cell Signaling Technology were: anti-phospho-SAMHD1-T592 (Cat# 15038), anti-SAMHD1 (Cat# 12361), and anti-CDC2/CDK1 (Cat# 9112). Mouse anti-β-actin MAb (Cat# 691001) was obtained from MP Biomedicals. Rat anti-hemagglutinin (HA) high-affinity antibody (Cat# 11-867-431-001) was obtained from Roche. Mouse anti-V5 and anti-V5-HRP antibodies (Cat# R960-25 and R961-25) were obtained from Invitrogen. Mouse anti-BGLF4 antibodies (Clones 2216 and 2224) were kindly provided by Mei-Ru Chen (Wang et al., 2005). High-sensitive anti-SAMHD1 antibodies (Cat# A311-354) were obtained from Bethyl Laboratories. Mouse anti-ZTA (Cat# 11-007) and anti-RTA (Cat# 11-008) antibodies were purchased from Argene. Mouse anti-BMRF1/EAD antibody was purchased from Millipore (Cat# MAB8186, clone R3).

### Phos-tag SDS-PAGE

The level of SAMHD1 phosphorylation was analyzed by applying samples to 6% (w/v) acrylamide SDS-PAGE gels supplemented with 50 μM Phos-tag and MnCl_2_ according to manufacturer’s instructions (Wako, Cat# AAL-107) (Kinoshita et al., 2006). This method allows simultaneous analysis of phosphoprotein isoforms and its non-phosphorylated counterpart in SDS-PAGE. The acrylamide-pendant Phos-tag ligand provides a phosphate affinity SDS-PAGE for mobility shift detection of phosphorylated proteins. When cell extracts were analyzed on Phos-tag acrylamide gel, cell extracts were prepared in 1x SDS-PAGE sample buffer without EDTA.

### Protein Expression and Purification

Halo-tagged WT and KD CHPKs proteins were expressed and purified as previously described (Li et al., 2015; Zhang et al., 2017). Briefly, Halo-tagged WT and KD CHPKs were transfected into 293T cells. Two T175 flasks of transfected cells were harvested 48 hrs post-transfection at 100% confluence and lysed with 25 ml HaloTag Protein Purification Buffer (50 mM HEPES pH7.5, 150 mM NaCl, 1mM DTT, 1mM EDTA and 0.005% NP40/IGEPAL CA-630) with Protease Inhibitor Cocktail. WT and KD Halo-CHPKs were enriched using the Halo-tag resin and CHPKs were eluted from the resin by washing 3 times with 0.5 ml HaloTag Protein Purification Buffer containing 20 μl Halo-TEV protease.

For SAMHD1 purification from *E. coli*, the pGEX-5x-2-SAMHD1 plasmid was transformed into competent *E.coli* BL21. The transformed bacterial were then cultured in LB media at 37 °C to OD600=0.8. Protein expression was induced in the presence of 0.1 mM IPTG at 16 °C overnight. The cells were harvested by centrifugation at 8000rpm for 10 min. The cell pellets were frozen and thawed two times and then re-suspended in PBS solution with 1 mM DTT and 0.1 mM PMSF. After sonication and centrifugation, the lysates were incubated overnight with prepared glutathione sepharose 4B (GE Healthcare, 17-0756-01). The beads were then washed and SAMHD1 protein was eluted with elution buffer (50 mM Tris-HCl, pH 7.5, 150mM NaCl, 5 mM MgCl_2_, 0.5 mM TCEP and 10% glycerol) containing Factor Xa overnight at 4 °C. The purified protein was stored at −80 °C.

### *In Vitro* Kinase Assay

Each sample was incubated in 40 μl Kinase Buffer containing 0.75 (v/v) magnesium-ATP cocktail buffer (Cat# 20-113; Upstate) and 6 μl of WT and KD CHPKs or CDK1/Cyclin B (Cat# 14-450, Upstate) for 30 min at 30°C. Reaction mixtures in Kinase Buffer without ATP were included as negative controls. Finally, reaction mixtures were separated by gel electrophoresis and phospho-SAMHD1 proteins were detected by Western Blot.

To generate phospho-SAMHD1 for dNTPase activity assay, SAMHD1 was *in vitro* phosphorylated using EBV BGLF4. CDK1/Cyclin B was used as a positive control. The kinase-dead BGLF4 and buffer were included as negative controls. In a phosphorylation reaction, full-length SAMHD1 was incubated at a ratio of 50:1 (w/w) with the kinases in 10 mM Tris-HCl pH 7.5, 150 mM NaCl, 5 mM MgCl2, 0.5 mM TCEP and 2mM ATP. The reaction was initiated by the addition of SAMHD1 and incubated for 6 hours at 4°C. The phosphorylation of SAMHD1 on T592 was confirmed by Western Blot (Arnold et al., 2015).

### *In vitro* dNTPase activity assay

All SAMHD1 dNTPase activity assays were performed in a reaction buffer (100 μl per reaction) containing 50 mM Tris-HCl, pH 7.5, 150 Mm NaCl, 5 mM MgCl2 and 0.5 mM TCEP. Recombinant SAMHD1 was phosphorylated by BGLF4 or CDK1 *in vitro*. Non-phosphorylated SAMHD1 was similarly processed by incubating with kinase-dead BGLF4 (BGLF4-KD) or Buffer control. Then phosphorylated and non-phosphorylated SAMHD1 (1 μM) were converted to GTP-bound dimer by adding 1mM GTP for 1 min. SAMHD1 dimer was further activated by adding dNTP mixture (12.5 μM each) for 1 min. The mixture was diluted 10-fold and the substrate dCTP, dTTP, dATP or dGTP (100 μM) is added for 0 to 240 min at room temperature. The reaction samples were collected at 0, 120 and 240 min and quenched by 10-fold dilution into ice cold buffer containing 10 mM EDTA, followed by spinning through an Amicon Ultra 0.5 ml 10 kDa filter at 16000 xg for 20 min (Jang et al., 2016; Tang et al., 2015).

Deproteinized samples were analyzed by HPLC using a Synergi C18 Column 150 × 4.6mm (Phenomenex). The column was pre-equilibrated in 20mM ammonium acetate, pH 4.5 (buffer A). Injected samples (100 μl) were eluted with a linear methanol gradient over 14 min at a flow rate of 1 mL/min. The dN yields were quantified by integration of the calibrated UV absorption peak at 260 nm.

### Lytic Induction

To induce the EBV lytic cycle in Akata B cells, Akata-BX1 (EBV+) and Akata (EBV+) cells were treated with IgG (1:200, Cat# 55087, MP Biomedicals) for 24, 48 and 72 hrs. Akata-4E3(EBV-) cells were treated similarly as controls. For lytic induction of EBV in P3HR-1 cells, the cells were treated by TPA (20 ng/ml; Cat# NC9325685, Fisher Scientific) and sodium butyrate (3 mM; Cat# 19-137, Millipore) for 24 and 48 hrs. For lytic induction of EBV in Hela-(EBV+) cells, the cells are transfected with EBV ZTA, RTA and ZTA plus RTA using Lipofectamine 2000 reagent (Cat# 11668019, Life Technologies) for 48 hrs. In some experiments, pHTPsiRNA-PK (siBGLF4) was co-transfected with ZTA/RTA to knockdown BGLF4.

### HCMV Infection

MRC-5 cells were either mock infected or infected at a multiplicity of infection (MOI) of 1 by HCMV virus TS15-rN, an epithelial-tropic variant of HCMV strain Towne (Cui et al., 2013) for 72 hrs. Stocks of virus TS15-rN were prepared and infectious titers determined as described previously (Cui et al., 2012).

### Virus Titration

EBV titers were determined using a Raji cell infection assay (Li et al., 2011; Meng et al., 2010). Briefly, GFP-EBV recombinant virus was harvested from lytically induced Akata BX1 cells carrying sgRNAs targeting SAMHD1 or non-targeting control sgRNA. Raji cells (2 × 10^5^ in 1 ml medium/well in 24-well plates) was infected with the GFP-virus and phorbol-12-myristate-13-acetate (TPA) (20 ng/ml) and sodium butyrate (3 mM) were added 24 h later. After a further 24hr, the GFP-positive Raji cells were scored using a fluorescence microscope. The number of green Raji cells was used to determine the concentration of infectious virus particles.

### EBV DNA Detection

To measure cell associated viral DNA, total genomic DNA was extracted using the Genomic DNA Purification Kit (Cat# A1120, Promega). The relative viral genome copy numbers were determined by quantitative PCR (qPCR) using primers to *BALF5* gene normalized by *β-actin.*

Primers sequences are: RL0062, 5’-agtccttcttggctagtctgttgac-3’ for BALF5-F; RL0063, 5’-ctttggcgcggatcctc-3’ for BALF5-R; RL0066, 5’-tggacttcgagcaagagatg-3’ for β-actin-F; RL0067, 5’-gaaggaaggctggaagagtg-3’ for β-actin-R.

### Statistical Analysis

Statistical analyses employed a two-tailed Student’s t test. A p value of ≤ 0.05 was considered statistically significant. Values are given as the mean of biological replicate experiments. Error bars represent the standard error of the mean or standard deviation from two or three biological replicates.

---

**Figure S1.**
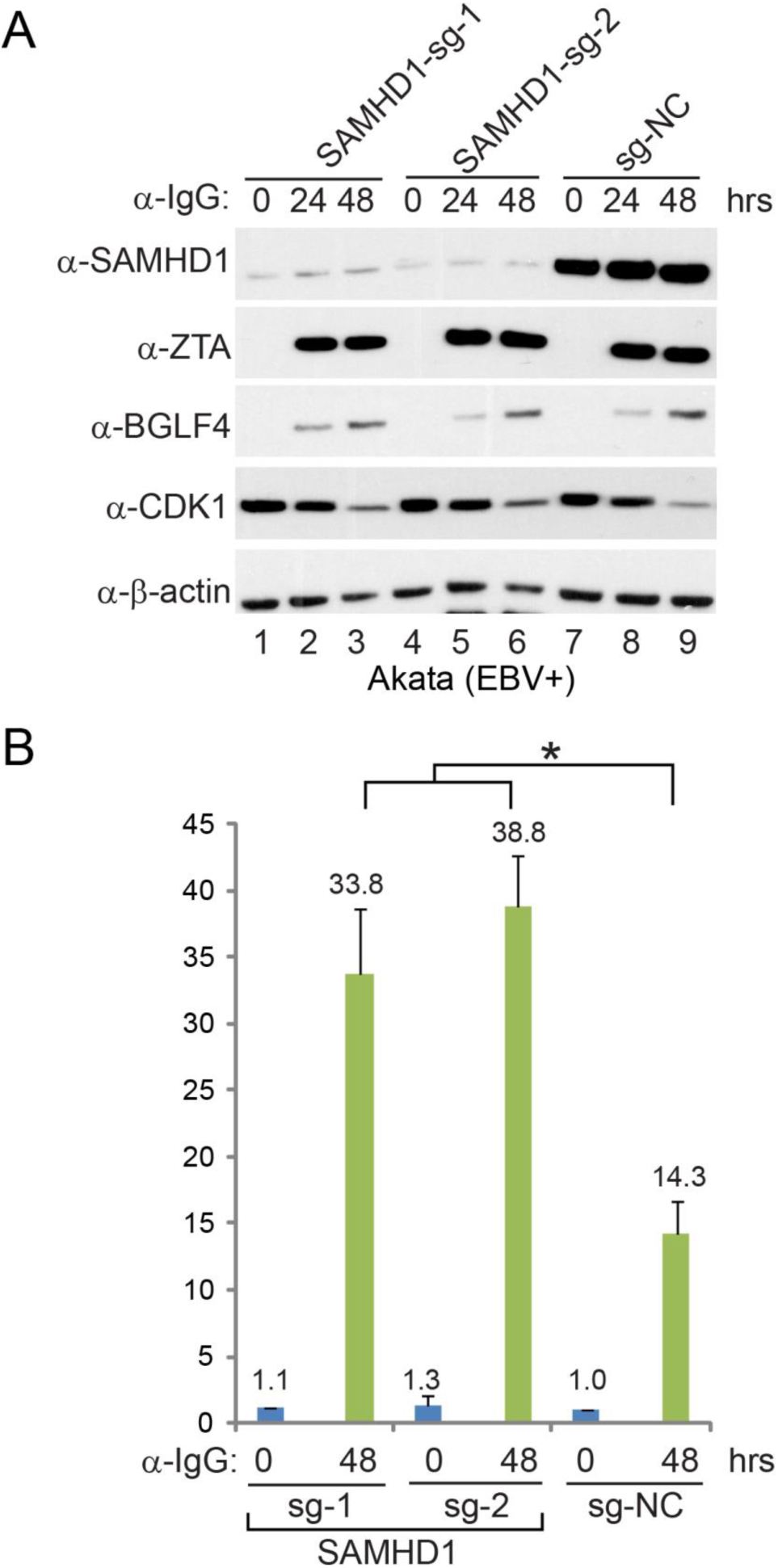
SAMHD1 depletion facilitates EBV lytic replication in Akata (EBV+) cells, related to Figure 3. (A) SAMHD1 depletion doesn’t affect EBV protein expression. SAMHD1-depleted and control Akata (EBV+) cells were treated with IgG cross-linking to induce EBV lytic reactivation. SAMHD1, the accumulation of EBV ZTA and BGLF4, and the CDK1 protein level were analyzed by immunoblotting at the indicated time points. β-actin served as a loading control (B) EBV DNA replication is enhanced in SAMHD1-depleted cells. DNA samples were extracted from Akata (EBV+) cells treated as indicated. The relative viral genome copy numbers were determined by quantitative PCR using primer to *BALF5* gene normalized by *β-actin*. Representative results from three biological replicates are presented. Error bars indicate the standard deviation. * p < 0.05.

**Figure S2.**
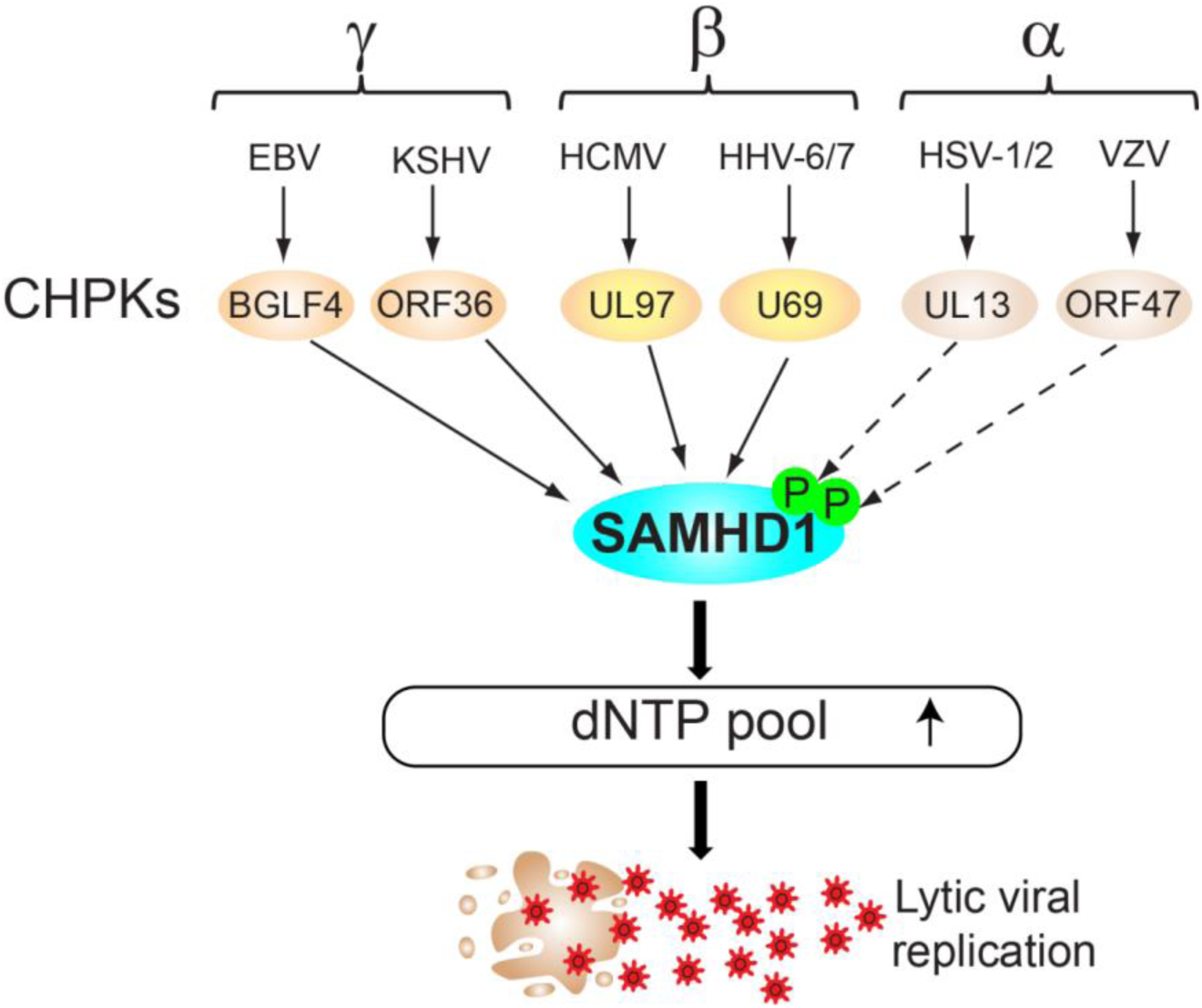
The model of SAMHD1 regulation by CHPKs in viral replication, related to Figures 6 and 7. Beta- and gamma-herpevirus protein kinases trigger the phosphorylation of SAMHD1, which leads to the increase of cellular dNTP pool for efficient viral DNA replication. The phosphorylation of SAMHD1 by beta- and gamma-herpevirus protein kinases is indicated by solid arrows. The possible regulation of SAMHD1 by alpha-herpevirus protein kinases is indicated by dashed arrows.

## Acknowledgements

We thank S. Diane Hayward (Johns Hopkins) for providing reagent and cells lines and for critically reading the manuscript. We thank Feng Zhang (MIT/Broad) for sharing the lentiCRISPR v2 plasmid (Addgene plasmid # 52961). We thank Didier Trono (EPFL) for providing the pMD2.G and psPAX2 plasmid (Addgene plasmid #s 12259 and 12260). We thank Robert Kalejta (University of Wisconsin-Madison) for providing the HA-tagged viral kinases (Addgene plasmid #s 26689-26698). We thank Michael McVoy and Ronzo Lee (Virginia Commonwealth University) for assistance with HCMV experiments. We thank Yong Xiong (Yale University) for providing HPLC column information. We thank Lindsey Hutt-Fletcher (Louisiana State University) for providing the Akata-BX1 (EBV+) cells and Mei-Ru Chen (National Taiwan University) for anti-BGLF4 antibody. We also thank Shannon Kenney (University of Wisconsin-Madison) for providing the Hela (EBV+) cell line and Edward Gershburg (Southern Illinois University) for providing siRNA-expressing plasmids.

## Author Contributions

RL and KZ conceived the project. RL, KZ, and DWL designed experiments. KZ, DWL and RL performed the experiments. RL and KZ wrote the manuscript.

## References

Arnold, L.H., Groom, H.C., Kunzelmann, S., Schwefel, D., Caswell, S.J., Ordonez, P., Mann, M.C., Rueschenbaum, S., Goldstone, D.C., Pennell, S., et al. (2015). Phospho-dependent Regulation of SAMHD1 Oligomerisation Couples Catalysis and Restriction. PLoS pathogens 11, e1005194.

Badia, R., Angulo, G., Riveira-Munoz, E., Pujantell, M., Puig, T., Ramirez, C., Torres-Torronteras, J., Marti, R., Pauls, E., Clotet, B., et al. (2015). Inhibition of herpes simplex virus type 1 by the CDK6 inhibitor PD-0332991 (palbociclib) through the control of SAMHD1. The Journal of antimicrobial chemotherapy.

Calderwood, M.A., Venkatesan, K., Xing, L., Chase, M.R., Vazquez, A., Holthaus, A.M., Ewence, A.E., Li, N., Hirozane-Kishikawa, T., Hill, D.E., et al. (2007). Epstein-Barr virus and virus human protein interaction maps. Proceedings of the National Academy of Sciences of the United States of America 104, 7606–7611.

Chang, C.W., Lee, C.P., Huang, Y.H., Yang, P.W., Wang, J.T., and Chen, M.R. (2012). Epstein-Barr virus protein kinase BGLF4 targets the nucleus through interaction with nucleoporins. Journal of virology 86, 8072–8085.

Chang, C.W., Lee, C.P., Su, M.T., Tsai, C.H., and Chen, M.R. (2015). BGLF4 kinase modulates the structure and transport preference of the nuclear pore complex to facilitate nuclear import of Epstein-Barr virus lytic proteins. Journal of virology 89, 1703–1718.

Chen, P.W., Lin, S.J., Tsai, S.C., Lin, J.H., Chen, M.R., Wang, J.T., Lee, C.P., and Tsai, C.H. (2010). Regulation of microtubule dynamics through phosphorylation on stathmin by Epstein-Barr virus kinase BGLF4. The Journal of biological chemistry 285, 10053–10063.

Chen, Z., Zhu, M., Pan, X., Zhu, Y., Yan, H., Jiang, T., Shen, Y., Dong, X., Zheng, N., Lu, J., et al. (2014). Inhibition of Hepatitis B virus replication by SAMHD1. Biochemical and biophysical research communications 450, 1462–1468.

Coquel, F., Silva, M.J., Techer, H., Zadorozhny, K., Sharma, S., Nieminuszczy, J., Mettling, C., Dardillac, E., Barthe, A., Schmitz, A.L., et al. (2018). SAMHD1 acts at stalled replication forks to prevent interferon induction. Nature 557, 57–61.

Cribier, A., Descours, B., Valadao, A.L., Laguette, N., and Benkirane, M. (2013). Phosphorylation of SAMHD1 by cyclin A2/CDK1 regulates its restriction activity toward HIV-1. Cell reports 3, 1036–1043.

Cui, X., Adler, S.P., Davison, A.J., Smith, L., Habib el, S.E., and McVoy, M.A. (2012). Bacterial artificial chromosome clones of viruses comprising the towne cytomegalovirus vaccine. Journal of biomedicine & biotechnology 2012, 428498.

Cui, X., Lee, R., Adler, S.P., and McVoy, M.A. (2013). Antibody inhibition of human cytomegalovirus spread in epithelial cell cultures. J Virol Methods 192, 44–50.

Daddacha, W., Koyen, A.E., Bastien, A.J., Head, P.E., Dhere, V.R., Nabeta, G.N., Connolly, E.C., Werner, E., Madden, M.Z., Daly, M.B., et al. (2017). SAMHD1 Promotes DNA End Resection to Facilitate DNA Repair by Homologous Recombination. Cell reports 20, 1921–1935.

Erazo, A., and Kinchington, P.R. (2010). Varicella-zoster virus open reading frame 66 protein kinase and its relationship to alphaherpesvirus US3 kinases. Current topics in microbiology and immunology 342, 79–98.

Feng, W.H., Kraus, R.J., Dickerson, S.J., Lim, H.J., Jones, R.J., Yu, X., Mertz, J.E., and Kenney, S.C. (2007). ZEB1 and c-Jun levels contribute to the establishment of highly lytic Epstein-Barr virus infection in gastric AGS cells. Journal of virology 81, 10113–10122.

Gershburg, E., Marschall, M., Hong, K., and Pagano, J.S. (2004). Expression and localization of the Epstein-Barr virus-encoded protein kinase. Journal of virology 78, 12140–12146.

Gershburg, E., and Pagano, J.S. (2008). Conserved herpesvirus protein kinases. Biochim Biophys Acta 1784, 203–212.

Gershburg, E., Raffa, S., Torrisi, M.R., and Pagano, J.S. (2007). Epstein-Barr virus-encoded protein kinase (BGLF4) is involved in production of infectious virus. Journal of virology 81, 5407–5412.

Gill, R.B., James, S.H., and Prichard, M.N. (2012). Human cytomegalovirus UL97 kinase alters the accumulation of CDK1. J Gen Virol 93, 1743–1755.

Hamirally, S., Kamil, J.P., Ndassa-Colday, Y.M., Lin, A.J., Jahng, W.J., Baek, M.C., Noton, S., Silva, L.A., Simpson-Holley, M., Knipe, D.M., et al. (2009). Viral mimicry of Cdc2/cyclin-dependent kinase 1 mediates disruption of nuclear lamina during human cytomegalovirus nuclear egress. PLoS pathogens 5, e1000275.

Hollenbaugh, J.A., Gee, P., Baker, J., Daly, M.B., Amie, S.M., Tate, J., Kasai, N., Kanemura, Y., Kim, D.H., Ward, B.M., et al. (2013). Host factor SAMHD1 restricts DNA viruses in non-dividing myeloid cells. PLoS pathogens 9, e1003481.

Hrecka, K., Hao, C., Gierszewska, M., Swanson, S.K., Kesik-Brodacka, M., Srivastava, S., Florens, L., Washburn, M.P., and Skowronski, J. (2011). Vpx relieves inhibition of HIV-1 infection of macrophages mediated by the SAMHD1 protein. Nature 474, 658–661.

Hume, A.J., Finkel, J.S., Kamil, J.P., Coen, D.M., Culbertson, M.R., and Kalejta, R.F. (2008). Phosphorylation of retinoblastoma protein by viral protein with cyclin-dependent kinase function. Science 320, 797–799.

Hwang, S., Kim, K.S., Flano, E., Wu, T.T., Tong, L.M., Park, A.N., Song, M.J., Sanchez, D.J., O’Connell, R.M., Cheng, G., et al. (2009). Conserved herpesviral kinase promotes viral persistence by inhibiting the IRF-3-mediated type I interferon response. Cell host & microbe 5, 166–178.

Iwahori, S., Hakki, M., Chou, S., and Kalejta, R.F. (2015). Molecular Determinants for the Inactivation of the Retinoblastoma Tumor Suppressor by the Viral Cyclin-dependent Kinase UL97. The Journal of biological chemistry 290, 19666–19680.

Iwahori, S., Murata, T., Kudoh, A., Sato, Y., Nakayama, S., Isomura, H., Kanda, T., and Tsurumi, T. (2009). Phosphorylation of p27Kip1 by Epstein-Barr virus protein kinase induces its degradation through SCFSkp2 ubiquitin ligase actions during viral lytic replication. The Journal of biological chemistry 284, 18923–18931.

Jang, S., Zhou, X., and Ahn, J. (2016). Substrate Specificity of SAMHD1 Triphosphohydrolase Activity Is Controlled by Deoxyribonucleoside Triphosphates and Phosphorylation at Thr592. Biochemistry 55, 5635–5646.

Kato, A., Yamamoto, M., Ohno, T., Tanaka, M., Sata, T., Nishiyama, Y., and Kawaguchi, Y. (2006). Herpes simplex virus 1-encoded protein kinase UL13 phosphorylates viral Us3 protein kinase and regulates nuclear localization of viral envelopment factors UL34 and UL31. Journal of virology 80, 1476–1486.

Kawaguchi, Y., and Kato, K. (2003). Protein kinases conserved in herpesviruses potentially share a function mimicking the cellular protein kinase cdc2. Reviews in medical virology 13, 331–340.

Kim, E.T., White, T.E., Brandariz-Nunez, A., Diaz-Griffero, F., and Weitzman, M.D. (2013). SAMHD1 restricts herpes simplex virus 1 in macrophages by limiting DNA replication. Journal of virology 87, 12949–12956.

Kinoshita, E., Kinoshita-Kikuta, E., Takiyama, K., and Koike, T. (2006). Phosphate-binding tag, a new tool to visualize phosphorylated proteins. Molecular & cellular proteomics : MCP 5, 749–757.

Kudoh, A., Daikoku, T., Ishimi, Y., Kawaguchi, Y., Shirata, N., Iwahori, S., Isomura, H., and Tsurumi, T. (2006). Phosphorylation of MCM4 at sites inactivating DNA helicase activity of the MCM4-MCM6-MCM7 complex during Epstein-Barr virus productive replication. Journal of virology 80, 10064–10072.

Kuny, C.V., Chinchilla, K., Culbertson, M.R., and Kalejta, R.F. (2010). Cyclin-dependent kinase-like function is shared by the beta-and gamma-subset of the conserved herpesvirus protein kinases. PLoS pathogens 6, e1001092.

Laguette, N., Sobhian, B., Casartelli, N., Ringeard, M., Chable-Bessia, C., Segeral, E., Yatim, A., Emiliani, S., Schwartz, O., and Benkirane, M. (2011). SAMHD1 is the dendritic-and myeloid-cell-specific HIV-1 restriction factor counteracted by Vpx. Nature 474, 654–657.

Lee, C.P., Chen, J.Y., Wang, J.T., Kimura, K., Takemoto, A., Lu, C.C., and Chen, M.R. (2007). Epstein-Barr virus BGLF4 kinase induces premature chromosome condensation through activation of condensin and topoisomerase II. Journal of virology 81, 5166–5180.

Lee, C.P., Huang, Y.H., Lin, S.F., Chang, Y., Chang, Y.H., Takada, K., and Chen, M.R. (2008). Epstein-Barr virus BGLF4 kinase induces disassembly of the nuclear lamina to facilitate virion production. Journal of virology 82, 11913–11926.

Lenzi, G.M., Domaoal, R.A., Kim, D.H., Schinazi, R.F., and Kim, B. (2015). Mechanistic and Kinetic Differences between Reverse Transcriptases of Vpx Coding and Non-coding Lentiviruses. The Journal of biological chemistry 290, 30078–30086.

Li, N., Zhang, W., and Cao, X. (2000). Identification of human homologue of mouse IFN-gamma induced protein from human dendritic cells. Immunology letters 74, 221–224.

Li, R., and Hayward, S.D. (2013). Potential of protein kinase inhibitors for treating herpesvirus-associated disease. Trends in microbiology 21, 286–295.

Li, R., Liao, G., Nirujogi, R.S., Pinto, S.M., Shaw, P.G., Huang, T.C., Wan, J., Qian, J., Gowda, H., Wu, X., et al. (2015). Phosphoproteomic Profiling Reveals Epstein-Barr Virus Protein Kinase Integration of DNA Damage Response and Mitotic Signaling. PLoS pathogens 11, e1005346.

Li, R., Wang, L., Liao, G., Guzzo, C.M., Matunis, M.J., Zhu, H., and Hayward, S.D. (2012). SUMO binding by the Epstein-Barr virus protein kinase BGLF4 is crucial for BGLF4 function. Journal of virology 86, 5412–5421.

Li, R., Zhu, J., Xie, Z., Liao, G., Liu, J., Chen, M.R., Hu, S., Woodard, C., Lin, J., Taverna, S.D., et al. (2011). Conserved herpesvirus kinases target the DNA damage response pathway and TIP60 histone acetyltransferase to promote virus replication. Cell host & microbe 10, 390–400.

Lv, D.W., Zhang, K., and Li, R. (2018). Interferon regulatory factor 8 regulates caspase-1 expression to facilitate Epstein-Barr virus reactivation in response to B cell receptor stimulation and chemical induction. PLoS pathogens 14, e1006868.

Ma, Z., Jacobs, S.R., West, J.A., Stopford, C., Zhang, Z., Davis, Z., Barber, G.N., Glaunsinger, B.A., Dittmer, D.P., and Damania, B. (2015). Modulation of the cGAS-STING DNA sensing pathway by gammaherpesviruses. Proceedings of the National Academy of Sciences of the United States of America 112, E4306–4315.

McGeoch, D.J., Cook, S., Dolan, A., Jamieson, F.E., and Telford, E.A. (1995). Molecular phylogeny and evolutionary timescale for the family of mammalian herpesviruses. Journal of molecular biology 247, 443–458.

Meng, Q., Hagemeier, S.R., Fingeroth, J.D., Gershburg, E., Pagano, J.S., and Kenney, S.C. (2010). The Epstein-Barr virus (EBV)-encoded protein kinase, EBV-PK, but not the thymidine kinase (EBV-TK), is required for ganciclovir and acyclovir inhibition of lytic viral production. Journal of virology 84, 4534–4542.

Moore, S.M., Cannon, J.S., Tanhehco, Y.C., Hamzeh, F.M., and Ambinder, R.F. (2001). Induction of Epstein-Barr virus kinases to sensitize tumor cells to nucleoside analogues. Antimicrobial agents and chemotherapy 45, 2082–2091.

Oberstein, A., Perlman, D.H., Shenk, T., and Terry, L.J. (2015). Human cytomegalovirus pUL97 kinase induces global changes in the infected cell phosphoproteome. Proteomics 15, 2006–2022.

Prichard, M.N. (2009). Function of human cytomegalovirus UL97 kinase in viral infection and its inhibition by maribavir. Reviews in medical virology 19, 215–229.

Prichard, M.N., Gao, N., Jairath, S., Mulamba, G., Krosky, P., Coen, D.M., Parker, B.O., and Pari, G.S. (1999). A recombinant human cytomegalovirus with a large deletion in UL97 has a severe replication deficiency. Journal of virology 73, 5663–5670.

Romaker, D., Schregel, V., Maurer, K., Auerochs, S., Marzi, A., Sticht, H., and Marschall, M. (2006). Analysis of the structure-activity relationship of four herpesviral UL97 subfamily protein kinases reveals partial but not full functional conservation. J Med Chem 49, 7044–7053.

Ruiz, A., Pauls, E., Badia, R., Torres-Torronteras, J., Riveira-Munoz, E., Clotet, B., Marti, R., Ballana, E., and Este, J.A. (2015). Cyclin D3-dependent control of the dNTP pool and HIV-1 replication in human macrophages. Cell cycle 14, 1657–1665.

Sanjana, N.E., Shalem, O., and Zhang, F. (2014). Improved vectors and genome-wide libraries for CRISPR screening. Nat Methods 11, 783–784.

Schwefel, D., Boucherit, V.C., Christodoulou, E., Walker, P.A., Stoye, J.P., Bishop, K.N., and Taylor, I.A. (2015). Molecular determinants for recognition of divergent SAMHD1 proteins by the lentiviral accessory protein Vpx. Cell host & microbe 17, 489–499.

Schwefel, D., Groom, H.C., Boucherit, V.C., Christodoulou, E., Walker, P.A., Stoye, J.P., Bishop, K.N., and Taylor, I.A. (2014). Structural basis of lentiviral subversion of a cellular protein degradation pathway. Nature 505, 234–238.

Sharma, M., Bender, B.J., Kamil, J.P., Lye, M.F., Pesola, J.M., Reim, N.I., Hogle, J.M., and Coen, D.M. (2015). Human cytomegalovirus UL97 phosphorylates the viral nuclear egress complex. Journal of virology 89, 523–534.

Sharma, M., and Coen, D.M. (2014). Comparison of effects of inhibitors of viral and cellular protein kinases on human cytomegalovirus disruption of nuclear lamina and nuclear egress. Journal of virology 88, 10982–10985.

Sharma, M., Kamil, J.P., Coughlin, M., Reim, N.I., and Coen, D.M. (2014). Human cytomegalovirus UL50 and UL53 recruit viral protein kinase UL97, not protein kinase C, for disruption of nuclear lamina and nuclear egress in infected cells. Journal of virology 88, 249–262.

Shi, Y., Lv, G., Chu, Z., Piao, L., Liu, X., Wang, T., Jiang, Y., and Zhang, P. (2014). Identification of natural splice variants of SAMHD1 in virus-infected HCC. Oncology reports 31, 687–692.

Shibaki, T., Suzutani, T., Yoshida, I., Ogasawara, M., and Azuma, M. (2001). Participation of type I interferon in the decreased virulence of the UL13 gene-deleted mutant of herpes simplex virus type 1. J Interferon Cytokine Res 21, 279–285.

Sullivan, V., Talarico, C.L., Stanat, S.C., Davis, M., Coen, D.M., and Biron, K.K. (1992). A protein kinase homologue controls phosphorylation of ganciclovir in human cytomegalovirus-infected cells. Nature 358, 162–164.

Sun, X., Bristol, J.A., Iwahori, S., Hagemeier, S.R., Meng, Q., Barlow, E.A., Fingeroth, J.D., Tarakanova, V.L., Kalejta, R.F., and Kenney, S.C. (2013). Hsp90 inhibitor 17-DMAG decreases expression of conserved herpesvirus protein kinases and reduces virus production in Epstein-Barr virus-infected cells. Journal of virology 87, 10126–10138.

Sze, A., Belgnaoui, S.M., Olagnier, D., Lin, R., Hiscott, J., and van Grevenynghe, J. (2013). Host restriction factor SAMHD1 limits human T cell leukemia virus type 1 infection of monocytes via STING-mediated apoptosis. Cell host & microbe 14, 422–434.

Tang, C., Ji, X., Wu, L., and Xiong, Y. (2015). Impaired dNTPase Activity of SAMHD1 by Phosphomimetic Mutation of Thr-592. The Journal of biological chemistry 290, 26352–26359.

Tarakanova, V.L., Leung-Pineda, V., Hwang, S., Yang, C.W., Matatall, K., Basson, M., Sun, R., Piwnica-Worms, H., Sleckman, B.P., and Virgin, H.W.t. (2007). Gamma-herpesvirus kinase actively initiates a DNA damage response by inducing phosphorylation of H2AX to foster viral replication. Cell host & microbe 1, 275–286.

Wang, J.T., Doong, S.L., Teng, S.C., Lee, C.P., Tsai, C.H., and Chen, M.R. (2009). Epstein-Barr virus BGLF4 kinase suppresses the interferon regulatory factor 3 signaling pathway. Journal of virology 83, 1856–1869.

Wang, J.T., Yang, P.W., Lee, C.P., Han, C.H., Tsai, C.H., and Chen, M.R. (2005). Detection of Epstein-Barr virus BGLF4 protein kinase in virus replication compartments and virus particles. J Gen Virol 86, 3215–3225.

Welbourn, S., Dutta, S.M., Semmes, O.J., and Strebel, K. (2013). Restriction of virus infection but not catalytic dNTPase activity is regulated by phosphorylation of SAMHD1. Journal of virology 87, 11516–11524.

Welbourn, S., Miyagi, E., White, T.E., Diaz-Griffero, F., and Strebel, K. (2012). Identification and characterization of naturally occurring splice variants of SAMHD1. Retrovirology 9, 86.

White, T.E., Brandariz-Nunez, A., Valle-Casuso, J.C., Amie, S., Nguyen, L.A., Kim, B., Tuzova, M., and Diaz-Griffero, F. (2013). The retroviral restriction ability of SAMHD1, but not its deoxynucleotide triphosphohydrolase activity, is regulated by phosphorylation. Cell host & microbe 13, 441–451.

Wittmann, S., Behrendt, R., Eissmann, K., Volkmann, B., Thomas, D., Ebert, T., Cribier, A., Benkirane, M., Hornung, V., Bouzas, N.F., et al. (2015). Phosphorylation of murine SAMHD1 regulates its antiretroviral activity. Retrovirology 12, 103.

Wolf, D.G., Courcelle, C.T., Prichard, M.N., and Mocarski, E.S. (2001). Distinct and separate roles for herpesvirus-conserved UL97 kinase in cytomegalovirus DNA synthesis and encapsidation. Proceedings of the National Academy of Sciences of the United States of America 98, 1895–1900.

Yan, J., Hao, C., DeLucia, M., Swanson, S., Florens, L., Washburn, M.P., Ahn, J., and Skowronski, J. (2015). CyclinA2-Cyclin-dependent Kinase Regulates SAMHD1 Protein Phosphohydrolase Domain. The Journal of biological chemistry 290, 13279–13292.

Zhang, K., Lv, D.W., and Li, R. (2017). B Cell Receptor Activation and Chemical Induction Trigger Caspase-Mediated Cleavage of PIAS1 to Facilitate Epstein-Barr Virus Reactivation. Cell reports 21, 3445–3457.

Zhu, J., Liao, G., Shan, L., Zhang, J., Chen, M.R., Hayward, G.S., Hayward, S.D., Desai, P., and Zhu, H. (2009). Protein array identification of substrates of the Epstein-Barr virus protein kinase BGLF4. Journal of virology 83, 5219–5231.

